# Seasonal patterns in *Synechococcus* pigment diversity at two temperate sites with contrasting oceanic regimes

**DOI:** 10.64898/2026.07.10.737521

**Authors:** Louison Dufour, Emile Faure, Frédéric Partensky, Francesco Mattei, Julia Uitz, Flavien Petit, Vincenzo Vellucci, Melek Golbol, Morgane Ratin, Bastian Gouriou, Martin Gachenot, Julia Clairet, Gregory K. Farrant, Mark Hoebeke, Erwan Corre, David Antoine, Anne-Claire Baudoux, Estelle Bigeard, Sarah Bureau, Jade Castel, Aurélie Chambouvet, Douglas Couet, Romain Cre’hriou, Colomban de Vargas, Céline Dimier, Florence Le Gall, Laure Guillou, Nicolas Henry, Fabienne Rigaut-Jalabert, Christian Jeanthon, Sarah Romac, Nathalie Simon, Jérémy Szymczak, Clara Trellu, Marie Walde, Anna Hickman, Stephanie Dutkiewicz, David M. Kehoe, Fabrice Not, Eric Thiébaut, Laurence Garczarek

## Abstract

Competition for light has driven extensive pigment diversification among phytoplankton species, yet how this diversity shapes their spatiotemporal distribution in the field has been little studied so far. The cyanobacterium *Synechococcus* is an ideal model for addressing this issue, since this group has colonized most light spectral niches in marine environments. Here, we used an approach based on marker read recruitment from metagenomes to analyze the seasonal succession of *Synechococcus* pigment types (PTs) at two time-series stations off French coasts exhibiting contrasting oceanic regimes. Marked seasonality was observed at both sites. The shallow, permanently mixed English Channel site SOMLIT-Astan was characterized by an alternation between green-light specialists (PT 3a) peaking in spring, and chromatic acclimaters type A (PT 3dA) accounting for most of the *Synechococcus* community in winter. In contrast, the pigment diversity was much higher at the deep Mediterranean station BOUSSOLE. In the upper layer, the two main PTs were the blue light specialists (PT 3c), which dominated the community in summer and fall, and PT 3dA cells, which were more abundant in spring. The third most abundant PT was chromatic acclimaters type B (PT 3dB), which accounted for up to 15% of the surface community in late fall. Strikingly, PT 3dA was dominant at depth during most of the year. Multivariate analyses between PT abundances, clade abundances and environmental factors, notably water color indexes, suggested new associations between PTs to specific clades and ecological niches. This study provides novel insights for refining distribution models of *Synechococcus* PTs and phytoplankton groups in general.

## INTRODUCTION

Light is required for photosynthesis, and competition for this critical resource has driven the extensive pigment diversity observed in modern phytoplankton species [1]. Understanding how this diversity influences the spatio-temporal dynamics of phytoplankton communities is crucial as climate change rapidly alters the underwater spectral light field [2–4]. *Synechococcus*, the second most abundant photosynthetic organism in the ocean, is an ideal biological model for exploring this question. Members of this group exhibit remarkable pigment and genetic diversity enabling their presence across a range of bio-optical regimes, from turbid estuaries to the clearest oligotrophic waters [5–7]. Like most other cyanobacteria, *Synechococcus* cells possess large, fan-shaped light-harvesting complexes called phycobilisomes. Phycobilisome rods are composed of different types of phycobiliproteins, which bind various types of chromophores (for more information see **Table S1**) [8–10]. Based on their phycobilisome composition, *Synechococcus* strains are divided into three main pigment types (PTs 1 to 3), independent from their vertical phylogeny [11, 12]. PT 3 is further divided into subtypes 3a through 3f, primarily based on the ratio of blue light-absorbing phycourobilin (PUB) chromophores to green light-absorbing phycoerythrobilin (PEB) chromophores [11–15]. While most pigment subtypes have a fixed PUB:PEB ratio, PT 3d strains can reversibly modify it to match the ambient light color (blue or green) via a process called type IV chromatic acclimation (CA4) [14, 16, 17]. CA4 ability is conferred by several genes present in a small genomic island that exists in two distinct configurations, CA4-A or CA4-B, and the corresponding strains are therefore called PTs 3dA and 3dB, respectively [14, 18–21] (**Table S1**).

Early studies on the biogeography of *Synechococcus* PTs used fluorescence-based methods such as epifluorescence microscopy, spectrofluorimetry, and/or flow cytometry [22–24]. With the development of molecular-based techniques, gene-based analyses targeting phycocyanin and/or phycoerythrin-I operons (i.e., *cpcBA* and *cpeBA*, respectively) have also been employed to study the geographical distribution of *Synechococcus* PTs [13, 25–32]. As in previous studies, PT 1 (i.e., phycocyanin-rich) cells were predominantly found in turbid, nutrient-rich, waters with low salinity, where red light penetrates the deepest. PT 2 cells were mainly encountered in eutrophic coastal areas, as well as in transition zones between brackish and marine environments where yellow-green light prevails. PT 3a cells (i.e., green specialists) predominated in onshore, green, mesotrophic waters. Cells assigned to the ‘PT 3c+3dB’ group were more abundant in warm oligotrophic waters, where blue light penetrates the deepest. Finally, PT 3dA cells were found in the temperate nutrient-rich waters of the English Channel [33] and constituted the whole *Synechococcus* community in the subpolar waters of the western Pacific Ocean [13]. Overall, these gene-based studies refined the spectral niches of the main *Synechococcus* PTs and subtypes, with the exception of PTs 3c and 3dB, which could not be distinguished using this approach [13, 33].

The combination of three additional phycobilisome markers, *cpcBA*, *mpeBA* and *mpeW,* allowed all known *Synechococcus* PTs to be distinguished from each other [14]. Extraction of these markers from the *Tara* Oceans metagenomic dataset enabled the establishment of the first global distribution map of *Synechococcus* PTs and subtypes, and highlighted the previously unknown global ecological significance of CA4-capable cells. Together, PTs 3dA and 3dB accounted for 41.5% of all *Synechococcus* reads among the sampled *Tara* Oceans stations. They were observed in complementary niches in the field with PT 3dA cells (22.6% of reads) being abundant in cold, nutrient-rich and highly productive waters, and PT 3dB cells (18.9% of reads) being primarily found in oligotrophic, iron-replete waters. Yet, both chromatic acclimaters appeared to be more abundant at depth. In addition, the study also refined the environmental parameters shaping the distribution of the two other major PT 3 subtypes. PT 3a cells (20.5% of reads) were associated with warm particle-rich waters and PT 3c cells (33.4% of reads) with warm, iron-replete regions. Surprisingly, although another blue light specialist (PT 3f) was quite rare along the *Tara* Oceans transect [15], it dominated the *Synechococcus* community at certain oligotrophic stations in the eastern Indian Ocean, particularly in the upper mixed layer [32].

While the global distribution of the different *Synechococcus* PTs has been well documented, it does not necessarily allow for a direct prediction of the seasonal PTs succession, which remains poorly characterized. Global surveys like *Tara* Oceans are indeed often temporally blind, providing snapshots that do not capture local-scale dynamics (e.g., seasonal changes in underwater light spectral quality, temperature and nutrients cycles, stratification and mixing). As a consequence, our ability to understand the mechanisms driving PT succession and to adequately represent their dynamics in biogeochemical and ecosystem models remains limited. To fill this gap, we conducted a two-year time series at two marine sites with contrasting oceanic and bio-optical regimes, i.e. SOMLIT-Astan in the English Channel and BOUSSOLE in the northwestern Mediterranean Sea. A metagenomic approach was used to describe the seasonal variations of both *Synechococcus* PTs and genotypes (i.e., clades), and to highlight the main environmental factors associated with their respective abundances, given that phycobilisome genes have shown to exhibit a distinct evolutionary history from vertically inherited core genes [11, 12] (**Table S2**). Our analyses revealed marked and distinct seasonal successions at both sites, with much higher PT and clade diversity at the Mediterranean site than at the English Channel station. We also demonstrated the predominance of one type of chromatic acclimater (PT 3dA) in winter at both stations, emphasizing the ecological importance of phenotypic plasticity for *Synechococcus* adaptation to changing environments.

## MATERIALS AND METHODS

### Sampling locations

The [SOMLIT](https://www.somlit.fr/en/) (“Service d’Observation en Milieu Littoral”)-Astan (a.k.a. SOMLIT-offshore [34]) station is located in the Western English Channel, 2.5 NM miles off Roscoff, France (48°46’N-3°58’W) [35], while the [BOUSSOLE](https://doi.org/10.18142/1) (“BOUée pour l’acquiSition d’une Série Optique à Long termE”) station is situated in the Ligurian Sea, 32 NM off Nice, France (43°22’N-7°54’E) [36]. Both are part of long-term time series sampled at regular intervals for hydrological, planktonic and/or bio-optical analyses. The two study sites are characterized by contrasting oceanic regimes. SOMLIT-Astan exhibits quasi-permanently tidally mixed waters and a maximum depth of 60 m [35, 37], while BOUSSOLE displays a strong seasonality of the mixed layer depth (MLD) and a depth of 2,440 m [36, 38]. Samples used in this study were collected over two consecutive years: i) twice a month in surface (2 m) at the SOMLIT-Astan station from October 2020 to October 2022 (**Dataset 1**), and ii) monthly at the BOUSSOLE station from August 2020 to August 2022 at four depths (**Dataset 2)**: near the surface (5 m), at an intermediate depth (generally 20 m), at the deep chlorophyll maximum (DCM; depth varying from 30 to 50 m) and below the DCM (60 m). The difference in sampling design reflects the contrasting regimes of the two study sites. At SOMLIT-Astan, the well-mixed coastal site with limited vertical structure, bimonthly surface sampling is supposed to provide an adequate representation of the water column, and sufficient temporal resolution to capture short-term variability and seasonal succession. At BOUSSOLE, the oligotrophic open-ocean site characterized by strong seasonal stratification, priority was given to monthly sampling at four depths to resolve the vertical structure of PT assemblages, at the expense of temporal resolution. Consequently, direct quantitative comparisons of short-term variability between the two sites should be interpreted cautiously, as differences may partly reflect sampling frequency rather than true ecological differences.

### Phytoplankton counts and pigment content

Samples of 1.5 mL seawater were fixed with a mix of glutaraldehyde (25%, Sigma-Aldrich) and pluronic acid (10%, Sigma-Aldrich), following SOMLIT protocols (https://www.somlit.fr/en/parameters-and-protocols/). After fixation in the dark for 15 min, samples were frozen in liquid nitrogen, then stored at -80°C. The abundance of marine picocyanobacteria (*Synechococcus* and *Prochlorococcus*) was determined by flow cytometry (**Datasets 1 and 2**) using a NovoCyte Advanteon flow cytometer (Agilent), as previously described [39].

Phytoplankton pigment contents were determined by High Performance Liquid Chromatography (HPLC). For this purpose, 2.27 L of seawater was vacuum filtered through 25 mm GF/F glass filters (Whatman), which were flash-frozen and stored at -80°C until analysis [40].

### Physico-chemical and optical parameters

*In-situ* measurements of seawater temperature and conductivity were collected at the SOMLIT-Astan and BOUSSOLE stations using Sea-Bird Scientific SBE19+ V2 and SBE 911+ CTD units respectively, also equipped with a chlorophyll *a* (Chl *a*) sensor (WETStar and Chelsea Aquatracka III, respectively). Fifteen additional parameters, including inorganic nutrients, particulate organic carbon (POC) and nitrogen (PON), were measured at the SOMLIT-Astan station as part of the SOMLIT national observation program (**Dataset 1;** https://www.somlit.fr/en/parameters-and-protocols/). These additional measurements were performed on the same day (and within three hours) as the metagenomics, HPLC, CTD, and optical parameters (see below). As concerns BOUSSOLE, samples for inorganic nutrients were collected from the same bottles as for phytoplankton counts and pigments, and analyzed following SOMLIT protocols (see above).

Radiometric profiles were acquired at both sites by deploying a free-fall multi-spectral radiometric profiler (Compact Optical Profiling System, C-OPS, Biospherical Instruments Inc.) equipped with 19 optical-filter microradiometers for measuring downward irradiance, upward nadir irradiance (BOUSSOLE) or radiance (SOMLIT-Astan), and above water downward irradiance. A minimum of three consecutive C-OPS profiles (from surface to ∼50 m at SOMLIT-Astan, and to ∼100 m at BOUSSOLE) were acquired at each station, with the in-water profiler at >30 m distance from the ship [41]. Radiometric profiles were corrected for possible light fluctuations from above water measurements. A second-degree local polynomial was fit to the radiometric profiles and used to extrapolate measurements to subsurface. Surface irradiance acquisitions were also fitted with the same function to remove the effect of ship’s roll, and the value at the beginning of the cast was used as surface reference irradiance (Es). Water leaving radiance (Lw) was obtained as the upward radiance just beneath the sea surface multiplied by a factor of 0.543 to account for the air-sea interface [42]. In the case of upward irradiance, values just beneath the sea surface were first transformed into radiance by applying look-up-tables from [43]. The remote sensing reflectance (Rrs) was calculated as the ratio of Lw to Es [44]. To characterize the light color available for *Synechococcus* cells, we computed quantitative indicators based on blue-to-green band ratios of remote sensing reflectance (Rrs_490:555_) and underwater downward irradiance (Irr_490:555_). These wavelengths were chosen among the 19 measured by the C-OPS for their closest proximity to PUB and PEB absorption maxima (490 and 545 nm, respectively). Continuous profiles of blue-to-green irradiance ratios were interpolated from observations using the *interp* function of the akima R package [45], and plotted using the ggplot2 package [46].

### Metagenomic sampling and analyses

A total of 134 metagenomes were produced for this study (**Datasets 3 and 4**). For each sample, 20 L of seawater were collected from the surface using a manual pump at SOMLIT-Astan, and a CTD-rosette device equipped with 12 L Niskin bottles at BOUSSOLE at four different depths: surface (Depth 1), 20 m (Depth 2), DCM (Depth 3) and 60 m (Depth 4). The bacterial size-fraction (i.e., 0.22-3 μm) was collected by filtration through 3 μm and 0.2 μm 142 mm polycarbonate filters (Millipore) mounted in series, following *Tara* Oceans protocols [47]. The filters were flash-frozen, then stored at -80°C. Simultaneous DNA and RNA extraction was performed by cryogenic grinding and nucleic acid extraction using NucleoSpin RNA kits (Macherey-Nagel) [48]. The extracted DNA was quantified using a ND-1000 spectrophotometer (Nanodrop) and a Qubit PicoGreen assay (Invitrogen). DNA integrity was checked on a Bioanalyzer DNA High Sensitivity chip (Agilent). Library preparations were performed on the Fasteris, Life Science Genesupport SA platform (Plan-Les-Ouates, Switzerland) using the TruSeq DNA Nano Prep kit (Illumina) after a size selection step by mechanical shearing of the DNA to obtain average insert sizes of 250 bp. The libraries were sequenced on Novaseq S4 (Illumina) in a 2 x 150 bp paired end mode to obtain about 160 million reads per sample and a high proportion (>90%) of merged overlapping read pairs. The obtained reads were trimmed of primers using cutadapt v2.8 [49], merged using flash v2.2.00 [50] and quality checked using the CLC Genomics Workbench v5.2.1 [51]. The mean number of merged, quality-checked reads per metagenome was of 170M for SOMLIT-Astan and 166M for BOUSSOLE. Accession numbers as well as detailed numbers of raw reads and reads after merging and quality checking are given in Datasets 3 and 4.

### Assessing the relative abundance of pigment types and genotypes

All merged metagenomic reads were mapped to a database of 256 reference picocyanobacteria genomes [52] using mmseqs search with a sensitivity of 7.5. All reads with mapping hits on any of the reference genomes were extracted and re-mapped on the same database complemented by 722 outgroup genomes from other cyanobacterial taxa, this time applying thresholds of 80% identity and 90% coverage. To assess the relative abundance of *Synechococcus* genotypes, reads corresponding to the high-resolution single-copy core marker *petB*, encoding cytochrome *b*_6_, were extracted and taxonomically assigned at the clade level as previously described [53] using a *petB* reference database containing 586 sequences, including 221 from 17 distinct *Synechococcus* clades (**Datasets 5 and 6**). This annotation was achieved using blast with thresholds of 80% identity, 90% coverage and an e-value threshold of 0.0001, leading to a total of 7,584 (respectively 42,496) annotated reads at SOMLIT-Astan (respectively BOUSSOLE). The abundances of reads per clade were then normalized by *petB* mean length and the sequencing depth of each sample, then transformed using the Hellinger transformation to study variations in clade abundance at the *Synechococcus* community level. A similar approach was used to quantify the relative abundance of *Synechococcus* PTs using the same sets of markers (*cpcBA*, *mpeBA* and *mpeW*) as in [15]. Again, the reads corresponding to one of the markers in the mapping against 978 genomes were assigned to each marker by Blast, based on their best-hits against specific databases for each marker (**Datasets 7 and 8**). The *cpcBA* reads were used to discriminate PTs 1, 2B, 2A or 3. Then, *mpeBA* reads allowed us to further split PT 3 into 3aA, 3f, 3dA and 3c/3dB (**Table S1**). The latter group was then separated using *mpeW*, a gene specific to PT 3dB. In total, 18,475 reads could be associated with a PT marker at SOMLIT-Astan, and 194,938 at BOUSSOLE. The reads were then summed by PT and normalized by the mean length of each marker and the sequencing depth of each sample. Finally, the abundance of each PT was computed as previously described [15], and transformed using the Hellinger transformation.

### Statistical analyses

#### Pre-processing of environmental metadata

A total of 50 physico-chemical and optical parameters were available for the whole time-series at SOMLIT-Astan, and 55 at BOUSSOLE. Variables with more than 50% missing values, as well as those correlated to another variable with a Pearson coefficient above 0.95, were manually removed (variables with the least NA were kept and in case of equal number of NAs the easiest variable to interpret was kept). The remaining variables had only 0.9% missing values at BOUSSOLE (43 variable) and 6.4% at SOMLIT-Astan (41 variables). These missing values were imputed using the classification and regression trees approach available in the mice R package [54]. In addition to these environmental data, two functions of the date of sampling representative of seasonality were added to each dataset. For each sample, the values of these functions were defined as

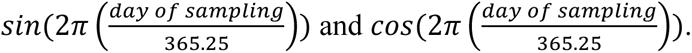

#### Permutations schemes for significance testing

To account for the time and depth autocorrelation structure of our sampling schemes, specific permutation designs were defined for each dataset. For SOMLIT-Astan, all statistical tests were based on permutations conserving chronology of the samples using the “series” type within the *how* command of the *permute* R package. Considering that we only had 50 samples over 2 years, we had only 100 distinct permutations available even after using the *mirror* option. To increase this number, and considering the strong annual reproducibility in our observed cycles (Fig. 1), the permutations were achieved independently on each year of sampling, from October to October, allowing to increase the maximum number of permutations from 100 to 10,000. At BOUSSOLE, the chronology of the samples was again conserved using the “series” type within the *how* command of the *permute* R package. To further take into account the fact that 4 discrete depths were sampled, permutations were achieved separately into 4 blocks, one for each depth, allowing to reach a maximum number of permutations >10^6^.

**Fig. 1:**
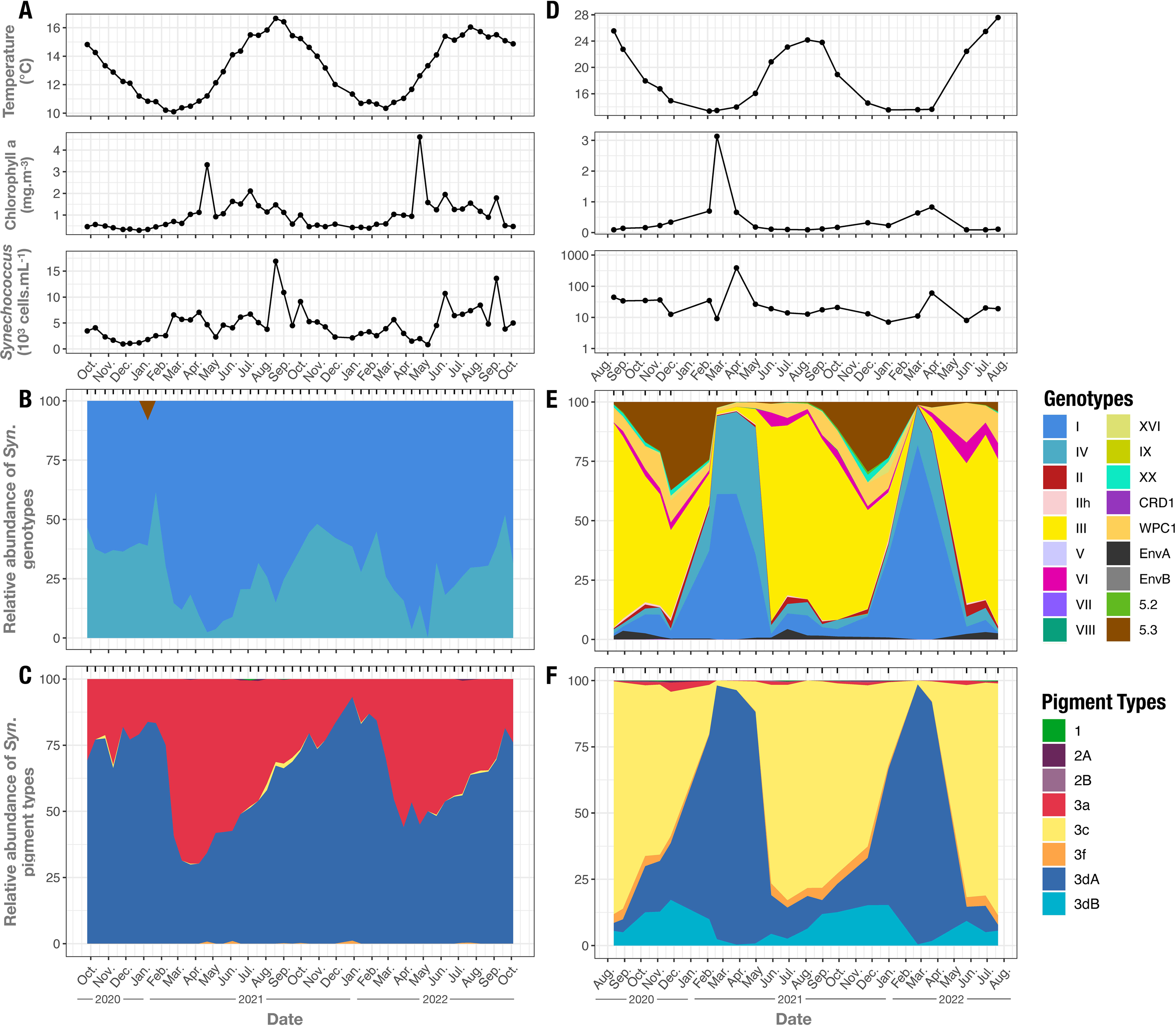
Seasonal variations of temperature, chlorophyll *a* concentration, *Synechococcus* cell abundance, genotypes and pigment types in surface waters at the SOMLIT-Astan station in the English Channel and the BOUSSOLE station in the Mediterranean Sea. **A**, **B**, **C** SOMLIT-Astan. **D, E, F** BOUSSOLE. **A, D** Temperature, chlorophyll *a* concentration, and *Synechococcus* cell abundance. **B, E** Relative abundances of *Synechococcus* genotypes at the clade level for members of subcluster (SC) 5.1 and at the SC level for SC 5.2 and 5.3. **C, F** Relative abundances of *Synechococcus* pigment types. Sampling points in B, C, E and F plots are indicated by black ticks at the top.

#### Univariate analysis

Spearman correlations between variables and Hellinger-transformed PT abundances in all samples were computed using the cor function in R, while p-values were calculated based on dataset-specific permutations (see above), and corrected using the Benjamini-Hochberg approach. All heatmaps were represented using the corrplot R package [56].

#### Multivariate analysis

To limit the overall collinearity across all potential predictors of PT abundance before conducting a multivariate analysis, we separated them into four categories: physico-chemistry, spatio-temporal structure (seasonality functions and depth for BOUSSOLE), genotypes and other biology (i.e., HPLC and flux cytometry measures). All the following variable and model selection steps are described in **Fig. S1**. Briefly, variables showing Pearson correlation coefficient above 0.7 or below -0.7 were removed within each category, again prioritizing easier to interpret variables (**Fig. S1**), except for genotypes data for which a PCA was fitted to obtain orthogonal descriptors of genotypic diversity without removing clades *a priori* (**Fig. S2**). A redundancy analysis model was then fitted on the selected variables or PCA axes using the *vegan* R package, again individually for each category. A stepwise model selection was performed on each full model to select the best parsimonious model in each category, using the *ordiR2step* R function in combination with our specific permutation schemes (see above). For each site, the four best models were then implemented in a variance partitioning analysis using the *varpart* R function. Finally, for visual purposes and based on the variance partition results, RDA models including both genotypes and physico-chemistry were built, after checking that the variance inflation factors (VIF) of each included variable remained below 10. In complement, a conditional RDA model investigating the effect of physico-chemistry alone at SOMLIT-Astan was built (including variables from all other categories as conditions), as well as one investigating the effect of genotypes alone at BOUSSOLE. Significance of all RDA models were assessed using our specific permutations schemes.

### Biogeochemical ecosystem model

We used the three-dimensional global Darwin model [57, 58] as a mechanistic framework to interpret the seasonal dynamics of *Synechococcus* PTs at BOUSSOLE only. Indeed, due to the coarse spatial resolution of the model, the shallow, tidally mixed, coastal SOMLIT-Astan station could not be directly represented. The model resolves major nutrient cycles and multiple plankton functional types, including the three main *Synechococcus* PTs (a representative green light specialist, blue light specialist and chromatic acclimater), [59], and explicitly represents size-dependent physiology and grazing, as well as spectrally resolved underwater light fields and phytoplankton absorption. Although designed for global biogeography rather than local variability, Darwin’s explicit treatment of light–mixing interactions and competition for light makes it well suited to test whether first-order seasonal PT patterns can be explained by physico–optical forcing alone.

## RESULTS AND DISCUSSION

### *Synechococcus* pigment types display marked seasonal dynamics in surface waters

Studying the seasonal successions of phytoplankton populations, a phenomenon particularly marked in temperate waters [60–62], is critical for refining our understanding of the factors that drive spectral niche partitioning. Here, we studied the variability of *Synechococcus* PTs and genotypes (i.e., clades) at two stations with contrasting environmental conditions: SOMLIT-Astan, a shallow, inshore station in the English Channel characterized by quasi-permanently mixed waters, and BOUSSOLE, a deep, seasonally stratified, offshore station of the Northwestern Mediterranean Sea.

Our measurements of *Synechococcus* cell abundance, environmental variables and pigment concentrations confirmed clear seasonal fluctuations of these parameters in surface waters at both stations (**Figs. 1A, 1D, S3 and S4**), consistent with previous reports in similar environments [36, 37, 63, 64]. At SOMLIT-Astan, the highest cell abundances (10^4^ cells mL^-1^) were observed between July and September each year (**Fig. 1A**), while a second bloom of slightly lower intensity (10^3^ cells mL^-1^) occurred in spring. At BOUSSOLE, a single *Synechococcus* bloom was observed each year in March-April (**Fig. 1D**). Maximum abundances of *Synechococcus* always coincided with the warmest seawater at SOMLIT-Astan (≈16°C) and with the coldest at BOUSSOLE (≈13°C; **Figs. 1A and 1D**). As expected, the variation of the concentration of zeaxanthin, a pigment specific to cyanobacteria [40], was fairly well synchronized with that of *Synechococcus* abundance (**Figs. 1A, 1D, S3C and S4A-B**). However, this was not the case for the temporal variations in Chl *a* concentration (**Figs. 1A and S4C**), confirming that eukaryotic phytoplankton dominated the Chl *a* biomass, especially during the spring bloom at SOMLIT-Astan [62].

At SOMLIT-Astan, metagenomic analyses revealed the year-round dominance of two PTs, green light specialist (PT 3a) and chromatic acclimater type A (PT 3dA; **Fig. 1C; Table S1)**, and two *Synechococcus* genotypes, clades I and IV (**Fig. 1B**). Such a low level of pigment and genetic diversity was previously reported for the same station in May 2012 [33]. The PTs displayed a very reproducible annual cycle and 60% of the variability in PT relative abundance could be explained by two simple seasonal sinusoid functions (**Fig. 2A**). Still, the best driver of PT abundance at this site was physico-chemistry, with 70% of the variance in PT abundance explained by only 4 variables (salinity, PO_4_^3-^, Irr_490:555_ and Rrs_490:555_; **Figs. S1A, 2A**). PT 3dA peaked in January, simultaneously with clade IV (**Fig. 1**) and in association with nutrient-rich, saline and cold waters (**Figs. 2, S5**). A sharp decrease of the 3dA:3a ratio in winter then led to a predominance of PT 3a in spring, matching the highest relative abundances of clade I (**Figs. 1B-C, 2**). We thus observe a strong similarity in the observed cycles of clades and PT relative abundances. All clade IV genomes sequenced to date are assigned to PT 3dA, while clade I genomes can be either PT 3a or 3dA [12, 15]. The strong correlation between PT 3a and clade I (**Fig. 2, S5**) suggests that PT 3a is abundant within clade I at this station. This assumption is supported by the observation that when the *Synechococcus* community consists almost entirely of clade I, PT 3a generally accounts for more than half of the cells (**Figs. 1B and 1C**). Interestingly, 10% of the variance in PT abundance could be explained by physico-chemistry only, even after removing potential effects from seasonality, genotype abundance and other biological parameters (p-value from the associated conditional RDA model = 0.005; **Fig**s. **2****, S6**). This result suggests that the within-clade PT variability observed at this site, i.e. the relative abundance of PT 3a and 3dA within clade I, can at least partly be explained by variations in salinity, nutrients and light quality. More precisely, the stronger presence of PT 3a in the late winter/early spring of 2021 compared to the same period in 2022 was associated with lower values of salinity, suggesting a potential impact of rainfall and runoff on the relative abundance of green-specialists at this coastal site (**Fig. S6**). This trend will have to be verified over a longer time-series. Very low levels of PT 3c were also observed in summer at SOMLIT-Astan, weakly linked to high Rrs_490:555_ values (**Fig. 2**), but these could not be associated with any specific genotype (**Fig. 1C**).

**Fig. 2:**
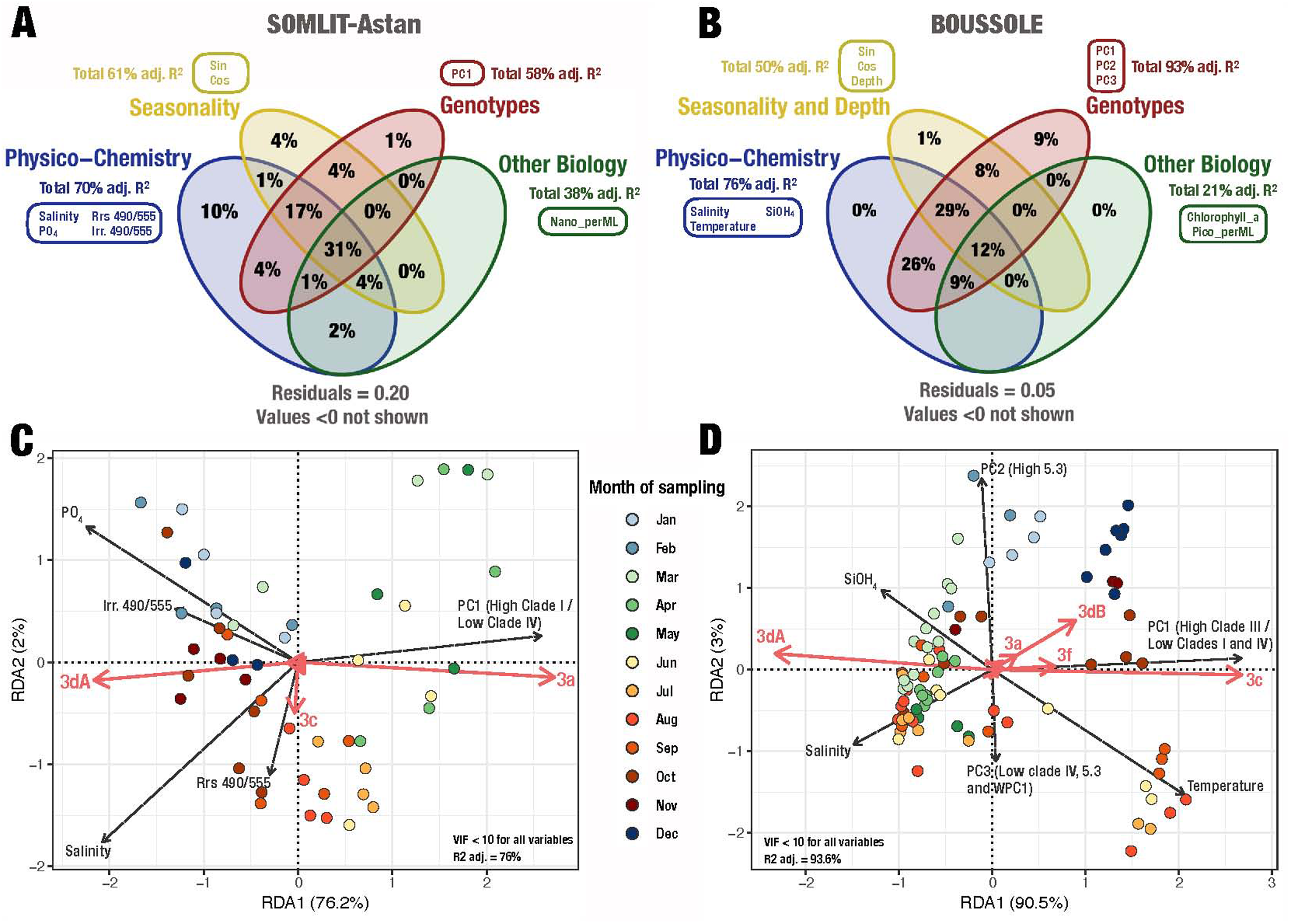
Multivariate analysis of the relationship between abundances of *Synechococcus* pigment types and potential predictors: physico-chemical data, seasonality, *Synechococcus* genotypes and other biological data. **A**,**B** Venn diagrams of the variance partition between 4 categories of predictors at SOMLIT-Astan (**A**) and BOUSSOLE (**B**), using Hellinger-transformed pigment type abundance as interest variable. The percentages indicated in the diagram reflect the amount of variance in pigment types abundance explained by the different combinations of predictors. For each category of predictors, the total sum of adjusted R^2^ is shown, as well as the list of selected variables (for precisions on the selection process, please refer to the Methods section and **Fig. S 1**). **C,D** Redundancy analysis (RDA) triplots for SOMLIT-Astan (**C**) and BOUSSOLE (**D**). Each RDA model was built using the selected physico-chemistry and genotypes variables as predictors of Hellinger-transformed pigment types abundance. The colored dots represent samples, colored according to their month of sampling. Dashed arrows represent the predictors, *i.e.*, physico-chemical variables and PCA axes reflecting genotypic diversity. For an easier interpretation, PCA axis were briefly described on the figure. PCA plots are available in **Fig. S2** . Plain arrows correspond to the different pigment types. Abbreviations: Irr. 490/555, ratio of downward irradiance at 490 nm and 555 nm; Rrs 490/55, ratio of remote sensing reflectance at 490 nm and 555 nm.

The BOUSSOLE site was also characterized by a reproducible PTs seasonal cycle but with much higher diversity than at SOMLIT-Astan, even though two PTs, blue-light specialist (3c) and chromatic acclimater type A (3dA), largely dominated the *Synechococcus* community at different periods of the year (**Fig. 1F**). This important diversity of PTs matched an increased diversity in genotypes compared to SOMLIT-Astan (**Figs. 1B, 1E**) and 93% of the variance in PTs abundances could be explained by the relative abundance of genotypes (compared to 58% in SOMLIT-Astan; **Fig. 2**). This suggests that there are virtually no within-genotype seasonal variations of PT at BOUSSOLE. PT 3dA predominated in surface waters in winter and spring, with maximum abundance in March (98% of the metagenomic reads in 2022), which coincided with the presence of clades I and IV (**Figs. 1E, 1F, 2 and S7)**. Unlike SOMLIT-Astan, most clade I members appeared to be PT 3dA, as more than 50% of the community was composed of clade I in Spring of each year, when PT 3dA accounted for >95% of the reads and PT 3a was scarce (<5%) (**Fig. 1E, 1F**). PT 3c was predominant during summer and fall, accounting for up to 88% of *Synechococcus* reads in August (**Fig. 1F**). This PT was strongly correlated with the major clade III (**Fig. 2, S7**), consistent with similar analyses carried out in September 2009 in the southwestern Mediterranean Sea (station TARA-009) [15, 53], but also with the minor clades EnvA and WPC1 (**Fig. S7)**. The third most abundant PT at BOUSSOLE was the chromatic acclimater type B (3dB), which accounted for up to 17% of the *Synechococcus* community in surface waters and exhibited its own seasonal pattern (**Fig. 1F**). Its relative abundance started increasing when the waters cooled down at the end of summer and peaked in late fall-early winter, matching the lowest values of salinity (**Fig. 2**), an observation missed by Grébert and coworkers [15] that helps refine the realized niche of this PT. Although PT 3dB is typically associated with strains (or genomes) from clades II, III and SC 5 [12, 14] (**Table S1**), the high similarity and correlation between the seasonal fluctuations of PT 3dB and SC 5.3 suggests that, at BOUSSOLE, this PT is mainly associated with SC 5.3 (**Figs. 1E, 1F, 2 and S7**). Similarly, the occurrence of another blue light specialist (PT 3f) appears to be restricted to a shorter period of the year (June to November), which coincides with the warmest part (>18°C) of the PT 3c temperature range (**Figs. 1D and 1F**). Surprisingly, even though the only two PT 3f strains (CC9616 and KORDI-100) characterized so far belong to clade XX [32], PT 3f relative abundance was not fully synchronized with that clade (**Figs. 1E and 1F**). This result may be attributed to the relatively high genetic diversity of PT 3f, which has been previously observed in the eastern Indian Ocean and suggested its occurrence in other clades [32]. Multivariate and correlation analyses suggested that PT 3f is associated with clade III at BOUSSOLE (**Figs. 2D** and **S7**), even though this association has not been observed so far in sequenced genomes. Altogether, these data highlight a seasonal succession of PTs at BOUSSOLE, with the blue-light specialists (PTs 3c and 3f) preferentially thriving in the warm oligotrophic waters (14-28°C and >18°C, respectively) of summer and fall, PT 3dB and the rare PT 3a occurring between late summer and late winter at lower salinity, and finally PT 3dA dominating in cold (<14°C), nutrient-rich waters in winter and spring. Our partition of variance analysis shows that no effect from physico-chemistry on PT abundances can be separated from variations in genotypes abundance at this offshore site (**Fig. 2**). Changes in the genotypes and PT diversity are thus inseparable at BOUSSOLE, while 17% of variance in PT abundance could be explained from physico-chemistry but not genotypes at SOMLIT-Astan. We further found that 9% of the variability in PTs abundances at BOUSSOLE could be explained by variations in genotype abundances even after removing potential effects from physico-chemistry, seasonality, depth and other biological parameters (p-value from the associated conditional RDA model < 0.001, **Fig. 2**). This share of variance thus corresponds to variations in genotypes that match variations in PTs, yet remains unexplained by contextual data. This could be linked with biotic competition between the highly diverse genotypes detected at this site, or to environmental factors that were not measured. Overall, our data allowed to explain 94% of the variations in PT abundance using only 6 variables (**Fig. 2**).

### Chromatic acclimaters type A dominate at depth throughout the year

In contrast to SOMLIT-Astan, where waters are mixed from the surface to the bottom throughout most of the year, the BOUSSOLE site is characterized by a marked stratification from May to September, typical of this particular oceanic regime [65], our rationale for analyzing depth profiles at this station. During this period, a sharp thermocline separated the warm (23-24°C), oligotrophic, Chl-poor, 15-20 m upper layer from the cold (13-14°C), nutrient-rich underlying water mass (**Dataset 2**). The water column was then mixed from November to April, resulting in homogeneous temperatures (**Fig. 3B**) and a gradual enrichment of the surface with inorganic nutrients (**Figs. 3D to 3F**).

**Fig. 3:**
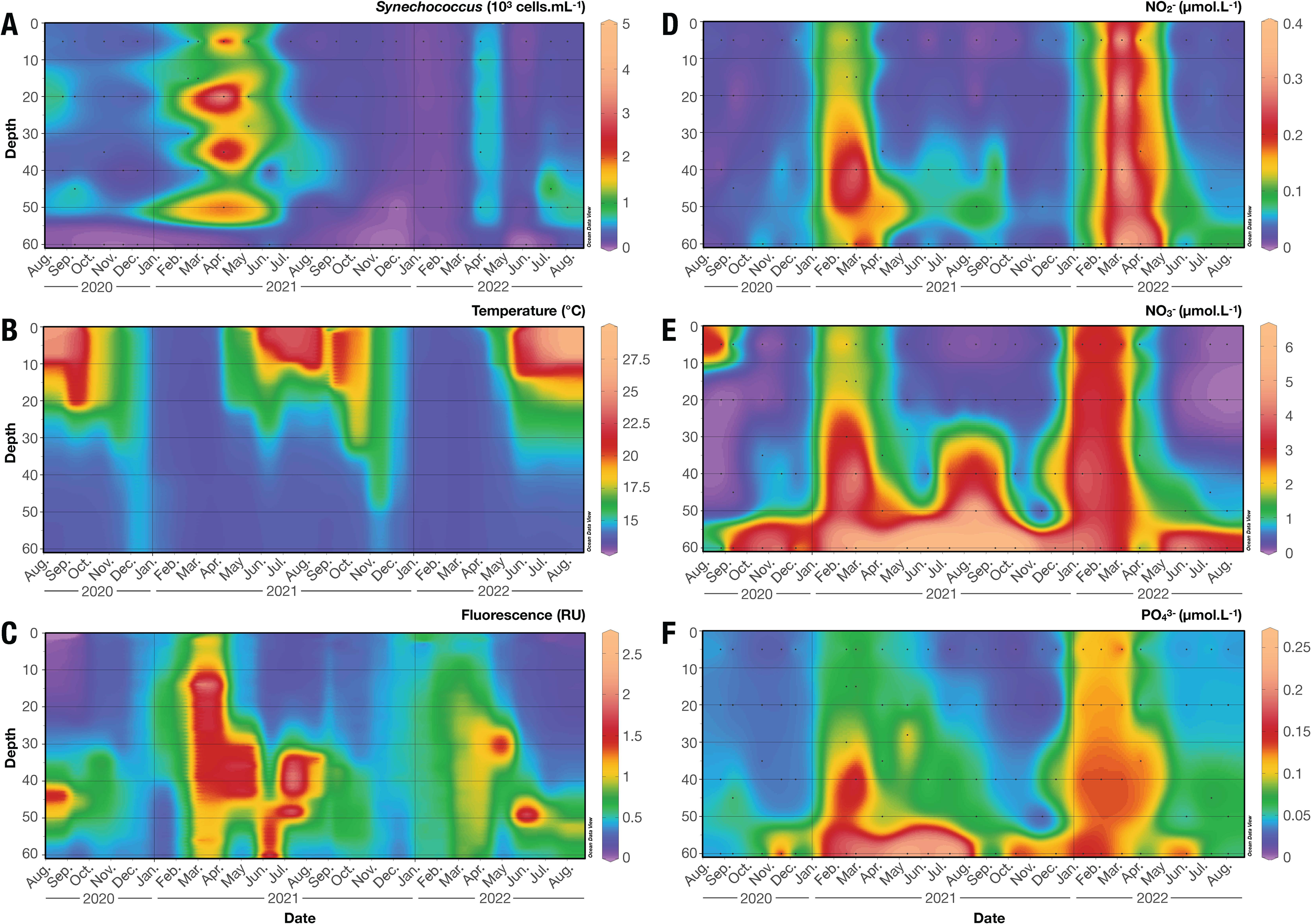
Seasonal variations across depths of *Synechococcus* cell abundance and physico-chemical parameters at BOUSSOLE. **A** *Synechococcus* cell abundance (10^3^ cells.mL^-1^). **B** Temperature (°C). **C** Fluorescence, a proxy of chlorophyll concentration (µg.L^-1^). **D**. Nitrite (NO_2_^-^) concentration (µmol.L^-1^). **E**. Nitrate (NO_3_^-^) concentration (µmol.L^-1^). **F**. Phosphate (PO_4_^3-^) concentration (µmol.L^-1^). Continuous profiles were obtained by interpolating the values between each individual observation, which are represented by black points.

Overall, the same PT diversity was observed at the surface and deeper in the water column at BOUSSOLE, with relative abundances varying strongly with depth. The same was true for genotypes, with the exception of a few minor clades (V and CRD1) that were restricted to the deepest samples (**Figs. 4A to 4F and S8A-B**). One of the most striking results of the present study is that the chromatic acclimater type A (PT 3dA) was by far the dominant PT at depth during most of the year, particularly at and below the DCM, where they accounted for 86 to 100% of the *Synechococcus* community from March to November (**Figs. 4A and S8B**). Consequently, the period during which blue light specialists (PTs 3c and 3f) dominated narrowed with depth, lasting from November to January at depth compared to May to January in surface for PT 3c (**Figs. 4B and S8A**). PTs 3c and 3f were not only significantly correlated with clade III, but also to several other minor clades such as VI, XX and WPC1 (**Figs. S7**). A similar narrowing trend was observed for the chromatic acclimater type B (PT 3dB, **Fig. 4C**). In contrast, the relative abundance of the green light specialists (PT 3a) appeared to be essentially invariant with depth (**Fig. S8B**). Altogether, the analysis of the seasonal variations in PTs abundances along the water column revealed the previously overlooked ecological importance of PT 3dA at depth in temperate waters, as well as novel PT/clade associations.

**Fig. 4:**
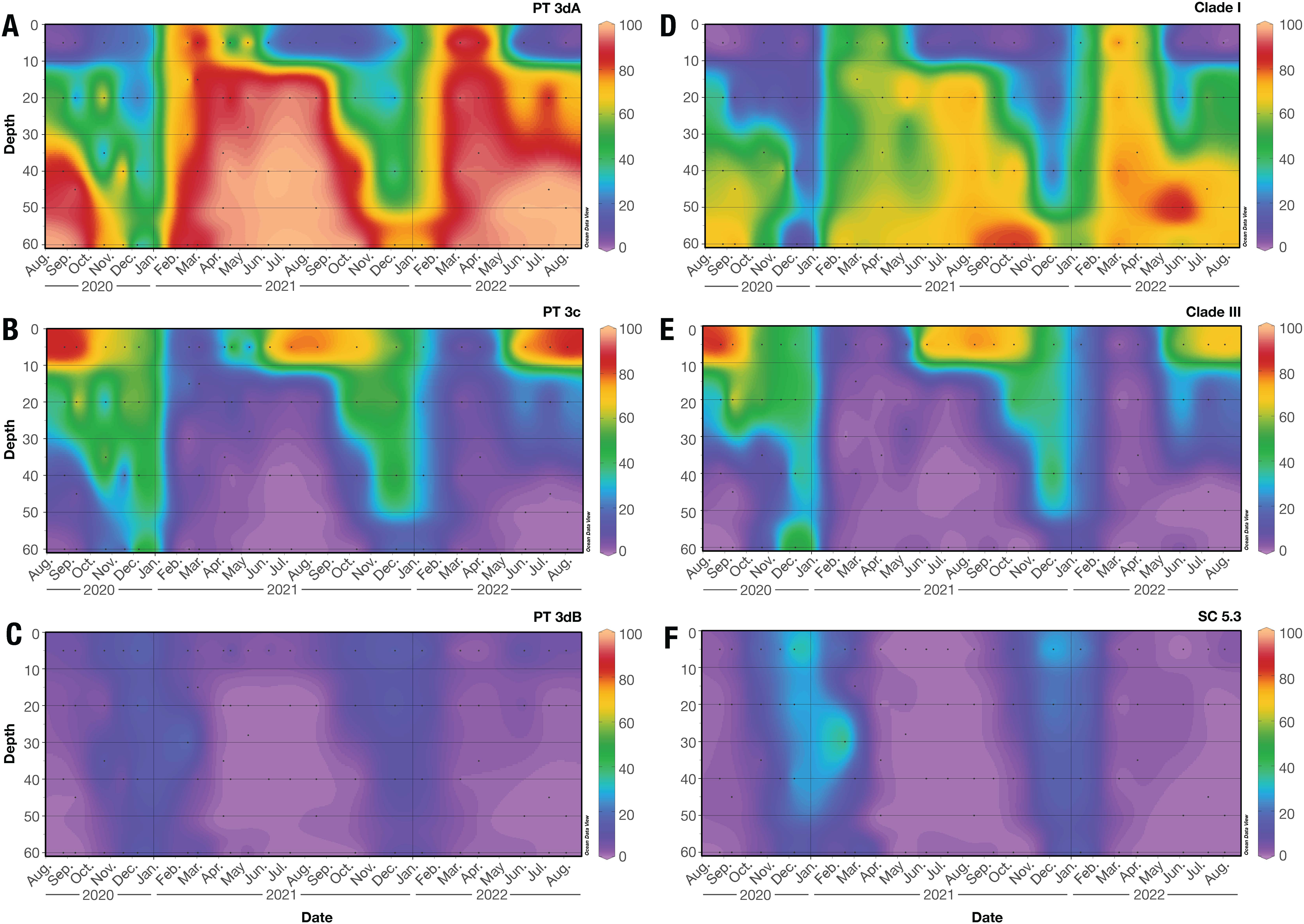
Seasonal variations of the relative abundances (%) of the main *Synechococcus* pigment types (PTs) and genotypes across depths at the Mediterranean station BOUSSOLE. **A** Chromatic acclimater type A (PT 3dA). **B** Blue light specialists (PT 3c). **C**. Chromatic acclimater type B (PT 3dB). **D**. Clade I. **E**. Clade III. **F**. Sub-cluster (SC) 5.3. These abundances were computed from metagenomic read recruitment on a set of reference gene markers (*cpcBA*, *mpeBA* and *mpeW* for PTs, *petB* for genotypes). Continuous profiles were obtained by interpolating the values between each individual observation, which are represented by black points. Note that, given the similar seasonal abundance patterns of clades I and IV, only the former is shown here. The relative abundances of all PTs and genotypes at the four sampling depths can be found in **Fig. S 8**.

### Effect of the underwater light field on seasonal and vertical variability of pigment types, and comparison with model data

To assess the role of the light field on the seasonal variations of *Synechococcus* PTs, data from the C-OPS profiler were used to retrieve a blue-to-green downward irradiance ratio (Irr_490:555_) at different depths, as well as a blue-to-green remote sensing reflectance ratio (Rrs_490:555_). As expected from the difference in trophic regimes between the two oceanic regions [5, 6, 66, 67], the coastal, chlorophyll- and nutrient-rich waters at SOMLIT-Astan were greenish throughout the year (Irr_490:555_ = 1.722 ± 0.704; Rrs_490:555_ = 1.250 ± 0.124; **Fig. 5A, Dataset 1**), while the warmer, oligotrophic waters at BOUSSOLE were predominantly bluish (Irr_490:555_ = 4.139 ± 3.142; Rrs_490:555_ = 2.050 ± 0.497; **Fig. 5B, Dataset 2**). Compared to SOMLIT-Astan, the Irr_490:555_ at BOUSSOLE was more variable and increased with depth, especially during periods of water mixing. The good correspondence between the vertical profiles of Chl *a* fluorescence (**Fig. 3C**) and Irr_490:555_ at BOUSSOLE (**Fig. 5B**) indicated that phytoplankton had a major effect in shaping the underwater spectral light field, as expected for Case I waters generally found at this site, i.e. open ocean waters for which optical properties are controlled by phytoplankton and their degradation products [36]. Despite showing only faint variations at SOMLIT-Astan (**Fig. 5A**), Irr_490:555_ and Rrs_490:555_ were both selected in our final RDA model at this site, with higher Irr_490:555_ values (bluer light) being associated with higher relative abundance of PT 3dA over the green light specialist PT 3a (**Figs. 2C and S6**).

**Fig. 5:**
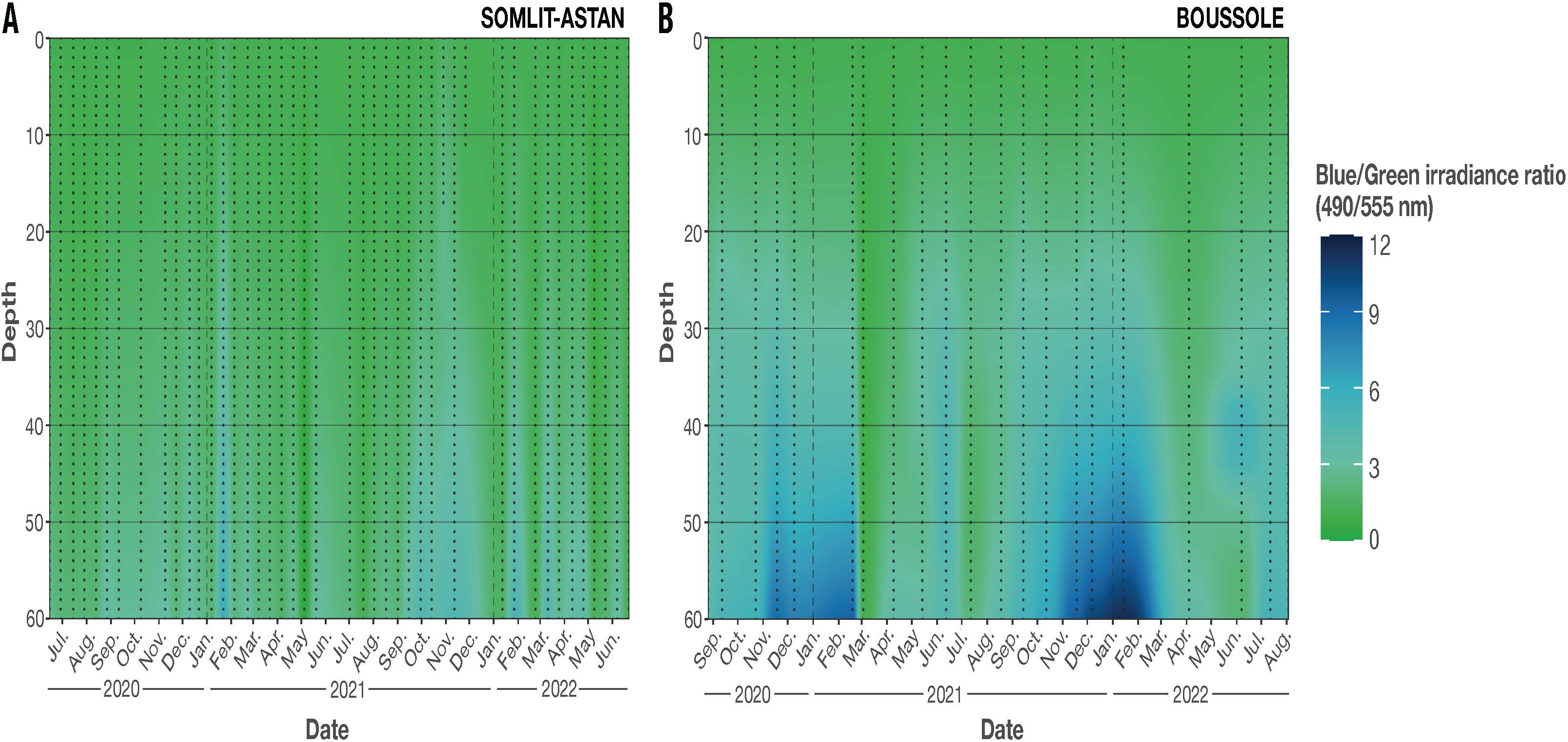
Seasonal variations across depths of the ratio of downward irradiance at 490 and 555 nm. **A** The English Channel station SOMLIT-Astan. **B** The Northwestern Mediterranean station BOUSSOLE. Continuous profiles were obtained by interpolating the values between each individual observation, which are represented by black points. The color scales were selected to reflect the overall underwater spectral field at both sites based on irradiance ratios.

The biological dataset obtained at the northwestern Mediterranean Sea BOUSSOLE time-series provides a critical observational framework for elucidating the seasonal dynamics of *Synechococcus* PTs and is particularly valuable in light of recent advances in modelling the distribution of PTs and their response to environmental factors [6, 59, 68]. Here, results from the Darwin model integrating the three main *Synechococcus* PTs correctly simulated the co-occurrence of blue-light specialists and chromatic acclimaters observed *in situ* at this site. However, the model failed to correctly assess their relative abundances (**Fig. S9**), and notably the predominance of chromatic acclimaters at depth (**Fig. 4A**). Both the model (**Fig. S10A**) and field measurements (**Fig. 5B**) indicated a predominantly blue underwater light field at depth at BOUSSOLE. The strong seasonality of PT relative abundances observed at this site, associated with alternation of stratified and mixed waters (**Figs. 3B and S8B**), is also consistent with the temporal dynamics of the corresponding PTs in the model (**Fig. S9**). Beyond reproducing these first-order patterns, the model also provides insights into the expected chromatic acclimaters’ acclimation states, used as a proxy for the PUB:PEB ratio exhibited by *Synechococcus* cells at any given season and depth (**Fig. S10B**). During periods of deep vertical mixing, *Synechococcus* cells experience a broader range of blue-to-green light conditions, resulting in the coexistence of multiple acclimation states in the modelled chromatic acclimater population (**Fig. S10B**). The capacity of chromatic acclimaters to exploit variability in the spectral light field has indeed been shown to confer them a competitive advantage [59, 68], consistent with their predominance during periods of strong spectral variability. Conversely, during months characterized by stronger stratification (May to August), cells experience a more stable spectral light field. In surface waters, where green wavelengths are more abundant, the majority of the modelled chromatically acclimating cells are in the greenest acclimation state (low PUB:PEB or CA1). In contrast, they converge toward the bluest acclimation state (high PUB:PEB or CA6) below the mixed layer, where the spectral field is blue-shifted (**Fig. S10B**).

Taken together, the abovementioned convergences between measured and modelled data at BOUSSOLE support the view that seasonal vertical mixing and its resulting spectral niche partitioning are strong drivers of PTs distribution. At the same time, the fine-scale seasonal patterns revealed by our time series, such as the dominance of PT 3dA at depth (**Fig. 4A**), while the model predicts a dominance of the blue light specialist (**Fig. S9**), indicate that light absorption properties alone are not always sufficient to explain the actual composition and dynamics of PT assemblages. These apparent contradictions underscore the importance of considering not only the PT of the major *Synechococcus* population, but also its genotype. Indeed, our statistical analysis of the BOUSSOLE metagenomics data suggests that all the PT diversity is directly linked with variations in genotypic diversity, which was not the case at SOMLIT-Astan. The cold-adapted clades I and IV are virtually the only clades present throughout the year at SOMLIT-Astan (**Fig. 1B**) and at depth at BOUSSOLE (**Fig. 4**), occupying a niche inaccessible to most other genotypes. As discussed earlier, clade I cells can be either chromatic acclimaters type A (3dA) or green light specialists (3a), while clade IV cells are exclusively 3dA [12]. Consequently, the observed dominance at depth of chromatic acclimaters type A instead of blue light specialists might not reflect a failure of optical niche theory but instead the superposition of optical selection with biogeographic and thermal constraints on genotype occurrence. Refining the representation of competition among dominant PTs will thus benefit from the integration of genotype-level traits, guided by seasonal *in situ* observations such as those presented here. In this context, discrepancies between model outputs and field observations should be viewed as valuable diagnostics to identify which ecological mechanisms, beyond optics, should be incorporated into the Darwin model to improve its predictive skills. This would enable us to better anticipate subtle, yet functionally meaningful, ecosystem responses to ongoing climate change.

## Conclusion

Seasonal fluctuations of environmental parameters have long been reported to be associated with dramatic changes in phytoplankton community structure and productivity throughout the year [37, 62, 69, 70]. Pigments have often been used as chemotaxonomic markers to identify taxonomic groups and assess the composition of marine phytoplankton communities, both *in situ* and from space [65, 71, 72]. However, few studies have so far analyzed the ecological significance of seasonal fluctuations of pigment assemblages, and especially annual successions of PTs within a given taxonomic group. Here, we focused on the abundant and ubiquitous cyanobacterium *Synechococcus,* a key component of phytoplankton communities. We showed that its extensive pigment diversity not only influences its wide spatial distribution in the world’s oceans [15], but also its seasonal occurrence. The present study also revealed that in temperate waters off French coasts, and likely temperate waters in general –generalizing beyond temperate systems will require dedicated time-series observations spanning a broader range of oceanographic conditions–, the predominant PT throughout most of the year is the chromatic acclimater type A (3dA). This pigment type alternated either with green light specialists (3a) in permanently mixed, greenish waters or with blue light specialists (3c) in seasonally stratified, surface waters. The dominance of PT 3dA cells at depth at BOUSSOLE is seemingly at odd with recent results from *in vivo* competition-for-light experiments, which showed that a blue light specialist won against a chromatic acclimater in low blue light, while a green specialist won against a chromatic acclimater in low green light [73]. This finding also differs from the results of *in silico* simulations with the Darwin model, which predicted that the blue light specialist should dominate at depth (**Fig. S9**). These apparent inconsistencies between culture-based experimental data, model predictions, and field observations highlight key ecological constraints that are not yet explicitly represented in current experimental and/or modeling frameworks. They underscore the importance of considering not only the PTs of the predominant *Synechococcus* population, but also their genotype (e.g., clades can be cold- or warm-adapted) as well as evolutionary relationships between PTs and genotypes since no clades encompass all PTs, and conversely a given PT may be found in different clades [11, 12] (**Table S2**). Such refinement would enable progress towards a more mechanistic representation of *Synechococcus* functional diversity in future ecosystem models, enhancing their predictive skill.

## DATA AVAILABILITY

The code used for metagenome analysis has been integrated into a snakemake workflow available on gitlab. Data, model outputs, and model configuration files generated during this study will be deposited in an open-access repository (e.g., Zenodo) upon acceptance of the manuscript for publication.

## ACKNOWLEDGMENTS

We warmly thank the crew of the R/V Neomysis and Tethys II for their help with sampling, the ABIMS platform for providing facilities for computational analysis and data storage, the SAPIGH platform for HPLC pigment analyses, Dominique Marie for technical support with flow cytometric analyses, Eric Macé for CTD analyses, and the IMEV staff (Eduardo Soto Garcia, Paco Still and Emilie Diamond) for hydrological data and nutrient analyses.

## AUTHORS CONTRIBUTION

DA, ET, FN, FPa, JU, LGa and VV contributed to funding acquisition. FPa, JU, and LGa conceived and supervised the project. AC, ACB, CD, CdV, CJ, CT, DC, EB, FPa, FPe, FRJ, JCa, JS, LD, LGa, LGu, MGo, MR, MW, NS, SB and SR and RC contributed to sampling for DNA and/or environmental parameters. LD and VV acquired and/or processed radiometric data. LD, LGa, CD, EF, JCl and MGa acquired and/or analyzed environmental data. BG, FLG, LD, JCa, MR and SR generated the metagenomes used in this study. EF and NH performed bioinformatic analyses. EC, FPa, GKF, LGa, MH and MR contributed to reference databases. EF performed the statistical analyses. FM performed modelling analyses. EF, LGa, FPa, LD, FM wrote the manuscript. AH, DA, DMK, EF, ET, FM, FPa, JU, LGa, LD, NS, SD and VV commented and edited the manuscript.

