## Supplemental Tables and Figures for "Seasonal patterns in *Synechococcus* pigment diversity at two temperate sites with contrasting oceanic regimes"

Deposited to BioRxiv: 2026-07-09

Keywords: Genomic observatory, marine cyanobacteria, metagenomics, molecular ecology, ocean color, phycobilisomes, phytoplankton, time series, niche adaptation

**Table S1: List of *Synechococcus* pigment types and corresponding phycobilisome rod phycobiliprotein and phycobilin composition.**
Abbreviations: n.a., not applicable; PCB, phycocyanobilin; PEB, phycoerythrobilin; PUB, phycourobilin.

| Pigment type | Phycocyanin | | Phycoerythrin-I | | Phycoerythrin-II | | PUB:PEB ratio | Color preferenda |
|---|---|---|---|---|---|---|---|---|
| | Localization | Phycobilin(s) | Localization | Phycobilin(s) | Localization | Phycobilin(s) | | |
| 1 | Entire rods | PCB | n.a. | n.a. | n.a. | n.a. | n.a. | Red light specialist |
| 2 | Basal rod disc | PCB | Other rod discs | PEB | n.a. | n.a. | n.a. | Yellow light specialist |
| 3a | Basal rod disc | PCB, PEB | Middle rod discs | PEB | Distal rod discs | PEB, PUB | Low (0.4) | Green light specialist |
| 3b | Basal rod disc | PCB, PEB and/or PUB | Middle rod discs | PEB, PUB | Distal rod discs | PEB, PUB | Intermediate | n.a. |
| 3c | Basal rod disc | PCB, PEB and/or PUB | Middle rod discs | PEB, PUB | Distal rod discs | PEB, PUB | High (1.6) | Blue light specialist |
| 3dA | Basal rod disc | PCB, PEB and/or PUB | Middle rod discs | PEB, PUB | Distal rod discs | PEB, PUB | Variable (0.6-1.6) | Chromatic acclimater A |
| 3dB | Basal rod disc | PCB, PEB and/or PUB | Middle rod discs | PEB, PUB | Distal rod discs | PEB, PUB | Variable (0.6-1.6) | Chromatic acclimater B |
| 3e | Basal rod disc | PCB, PEB and/or PUB | Middle rod discs | PEB, PUB | Distal rod discs | PEB, PUB | Variable (0.6-0.8) | n.a. |
| 3f | Basal rod disc | PCB, PEB and/or PUB | Middle rod discs | PEB, PUB | Distal rod discs | PEB, PUB | High (1.6) | Blue light specialist |

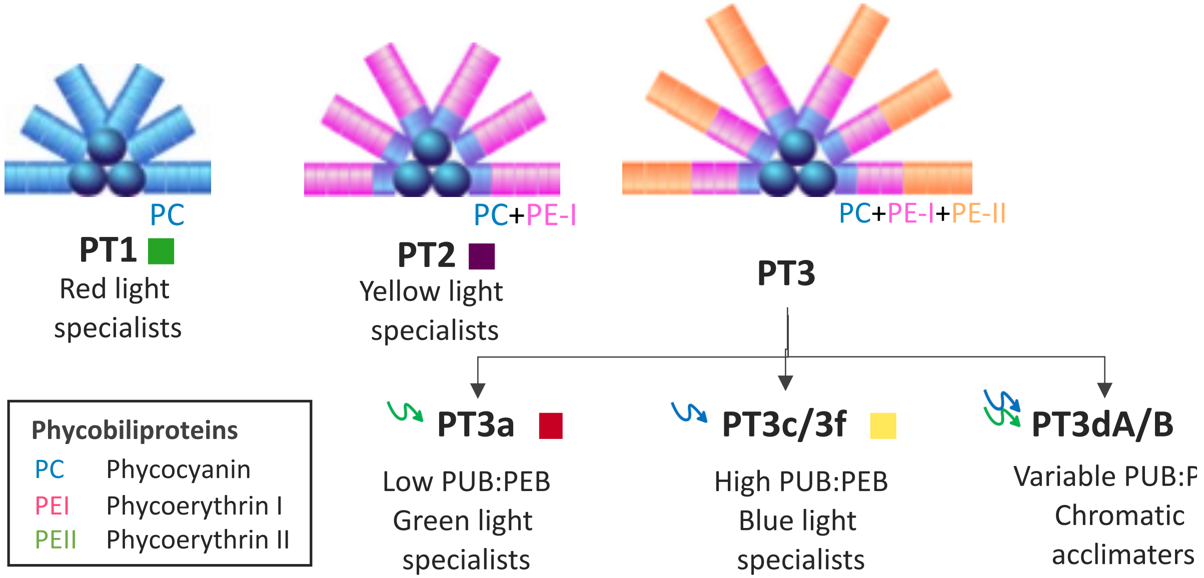

**Table S2: List of *Synechococcus* taxa and associated pigment types known in sequenced genomes so far.**

| Subcluster | Clade | Pigment types |
|---|---|---|
| 5.1 | I | 3a, 3dA, 3aA |
| 5.1 | II | 2, 3a, 3c, 3dB |
| 5.1 | III | 3c, 3dB |
| 5.1 | IV | 3dA |
| 5.1 | V | 2, 3a |
| 5.1 | VI | 2, 3c |
| 5.1 | VII | 3a, 3c |
| 5.1 | VIII | 1, 2 |
| 5.1 | IX | 3dA |
| 5.1 | XVI | 3dA |
| 5.1 | XX/UC-A | 3f |
| 5.1 | CRD1 | 3dA, 3cA |
| 5.1 | EnvA | 3c, 3f |
| 5.1 | EnvB | 3c |
| 5.1 | WPC1 | 3a, 3c |
| 5.2 | - | 1, 2, 3a |
| 5.3 | - | 2, 3dA, 3dB, 3eA |

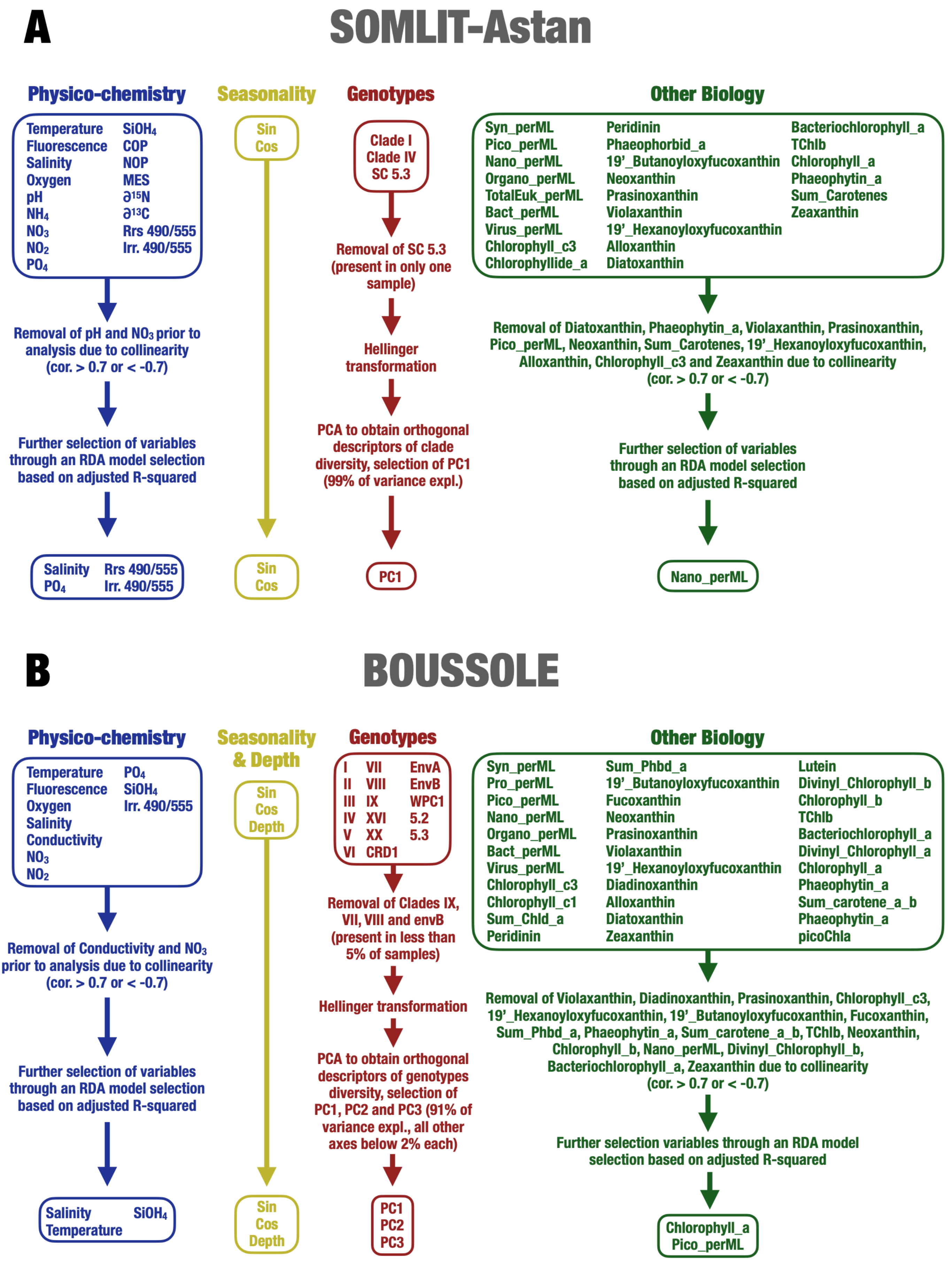


Fig. S1: Description of the variable selection process for multivariate analysis at SOMLIT (A) and BOUSSOLE (B).

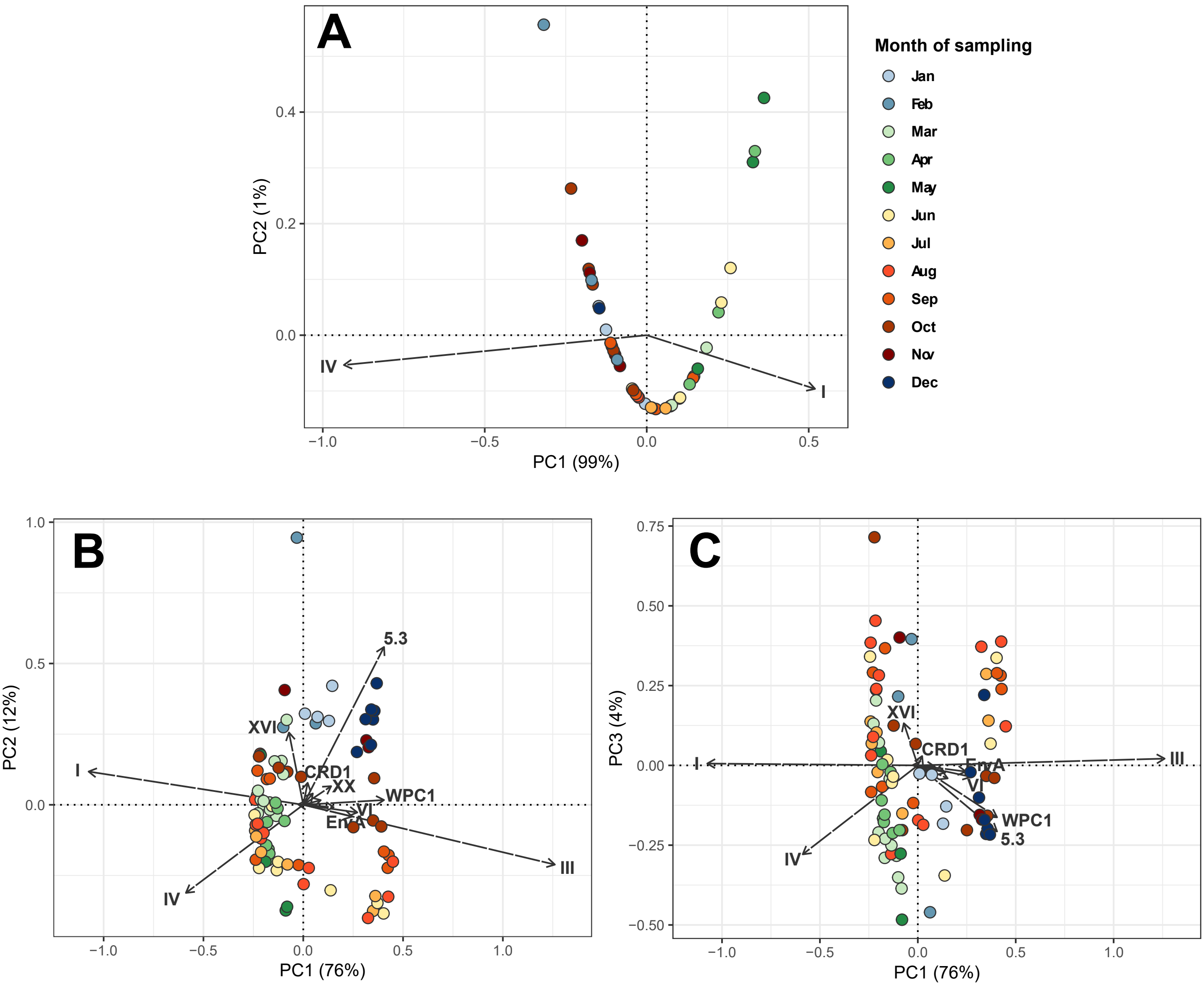


**Fig. S2: PCA of *Synechococcus* genotypes abundance at SOMLIT-Astan (A) and BOUSSOLE (B,C).** Genotype abundances as estimated from petB sequence abundance, normalized by sequencing depth and gene length before transformation with the Hellinger method. The coordinates of samples in PCA space were used as input in our redundancy analysis (RDA) models: PC1 only for SOMLIT-Astan (**A**), PC1, PC2 (**B**) and PC3 (**C**) for BOUSSOLE (See Figure SX and methods for full details on RDA models construction).

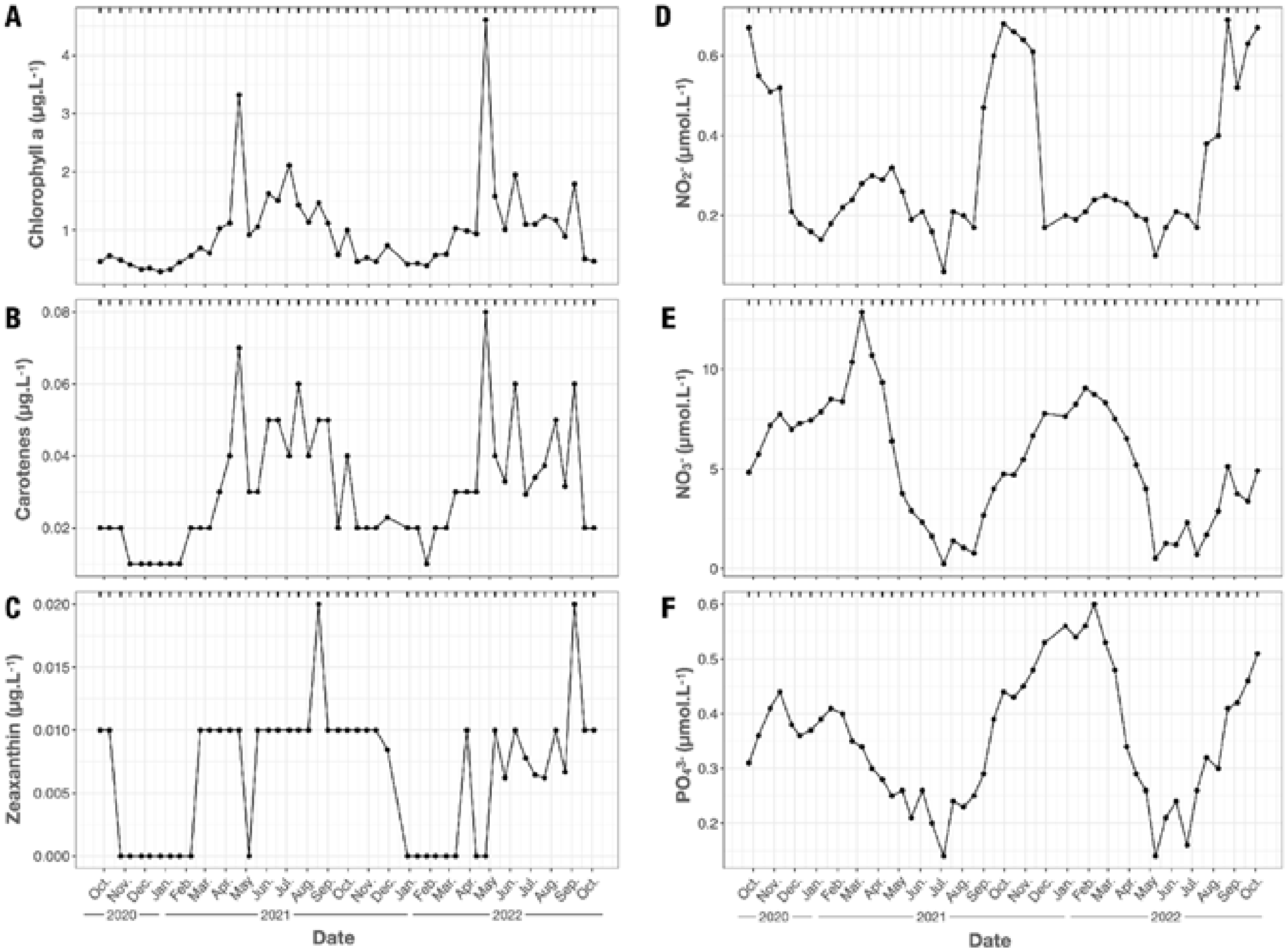


**Fig. S3: Seasonal variations of liposoluble pigments and inorganic nutrient concentrations at the English Channel station SOMLIT-Astan. A** Chlorophyll *a.* **B** Total carotenes. **C** Zeaxanthin. **D** Nitrite ($NO_2^-$). **E** Nitrate ($NO_3^-$). **F** Phosphate ($PO_4^{3-}$). The black ticks on top of each graph indicate the sampling dates.

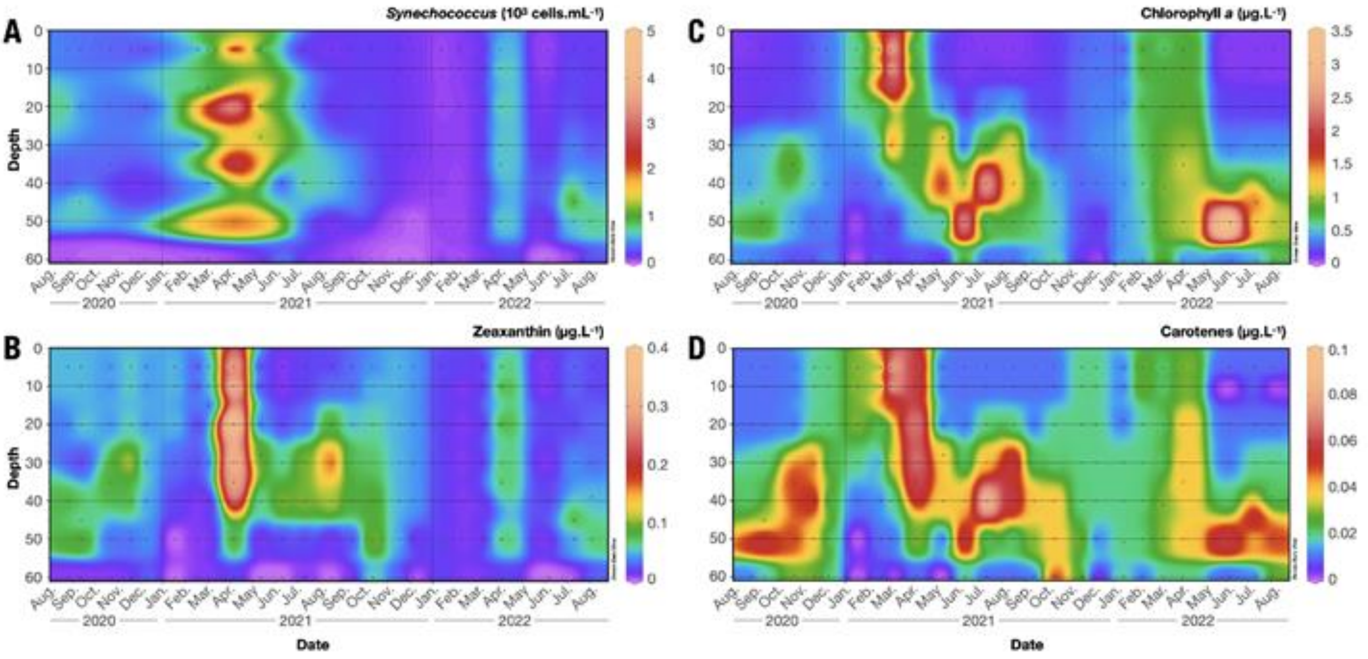


**Fig. S4: Seasonal variations across depths of liposoluble pigments as compared to *Synechococcus* abundance at the Mediterranean station BOUSSOLE. A** *Synechococcus* cell concentration. **B.** Zeaxanthin. **C.** Chlorophyll *a.* **D** Total carotenes. Continuous profiles were obtained by interpolating values between each individual observation, which are represented by black points.

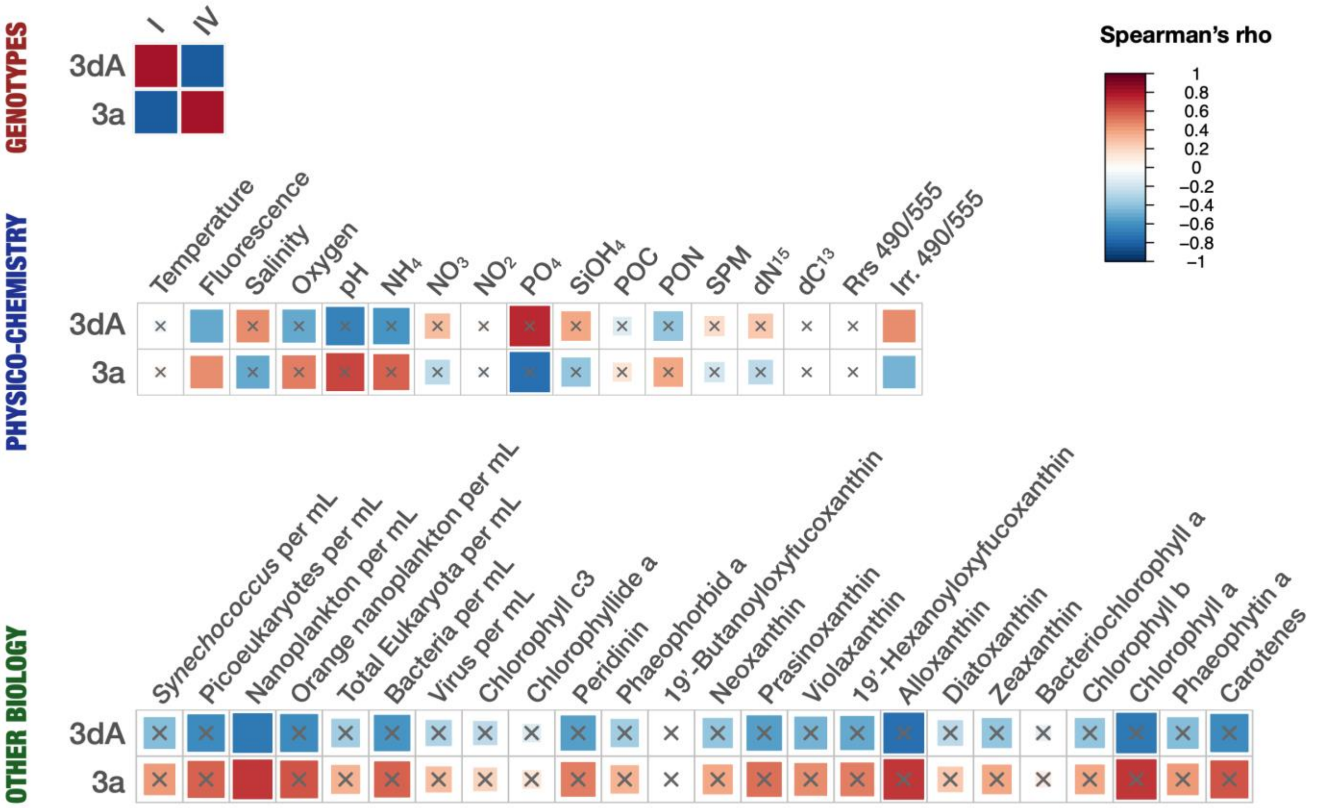


**Fig. S5: Correlation analysis between *Synechococcus* pigment types and all environmental and biological parameters measured at the English Channel station SOMLIT-Astan.** Each row corresponds to a pigment type and each column to a parameter, separated in the three categories: *Synechococcus* genotypes, physico-chemical parameters and other biological parameters. The scale indicates Spearman correlation coefficient between pairs of variables. Grey crosses indicate non-significant correlations (permutation-based test using 999 permutations, adjusted p-values > 0.05). Only pigment types and genotypes representing more than 5% of the *Synechococcus* reads in the whole dataset were represented. Abbreviations: POC, particulate organic carbon; PON, particulate organic nitrogen; SPM, suspended particulate matter; Irr. 490/555, ratio of downward irradiance at 490 nm and 555 nm; Rrs. 490/555, ratio of remote sensing reflectance at 490 nm and 555 nm. See Dataset 1 for more details.

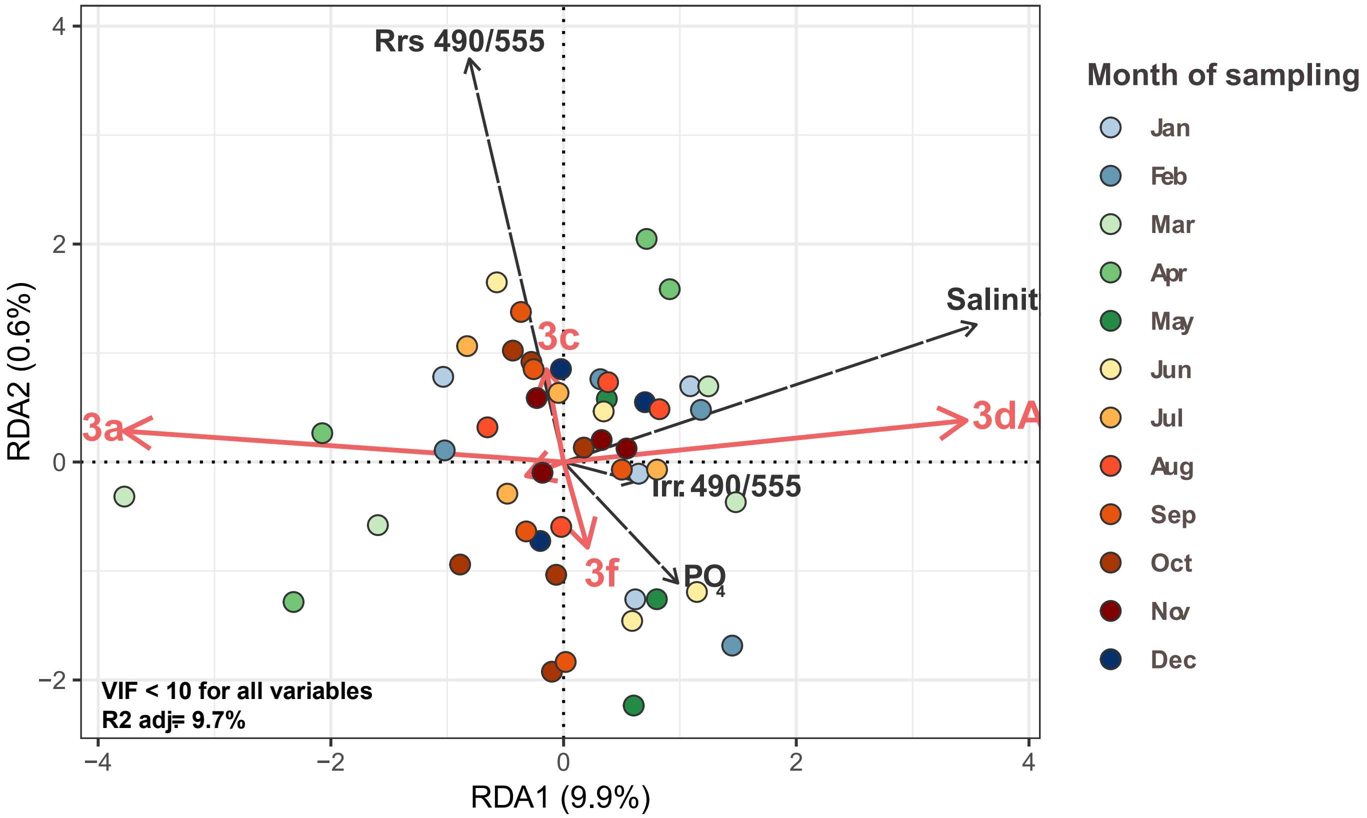


**Fig. S6: Conditional redundancy analysis (RDA) of the relationship between pigment types abundance and the physico-chemical context at SOMLIT-Astan.** Effects of seasonality (Sin and Cos), genotypes (PC1) and other biological factors (Nanoplankton per mL) were all included as conditioning variables in the RDA model. This way, this triplot reflects the impact of physico-chemistry on pigment types abundance after removing any covariation effects from these variables.

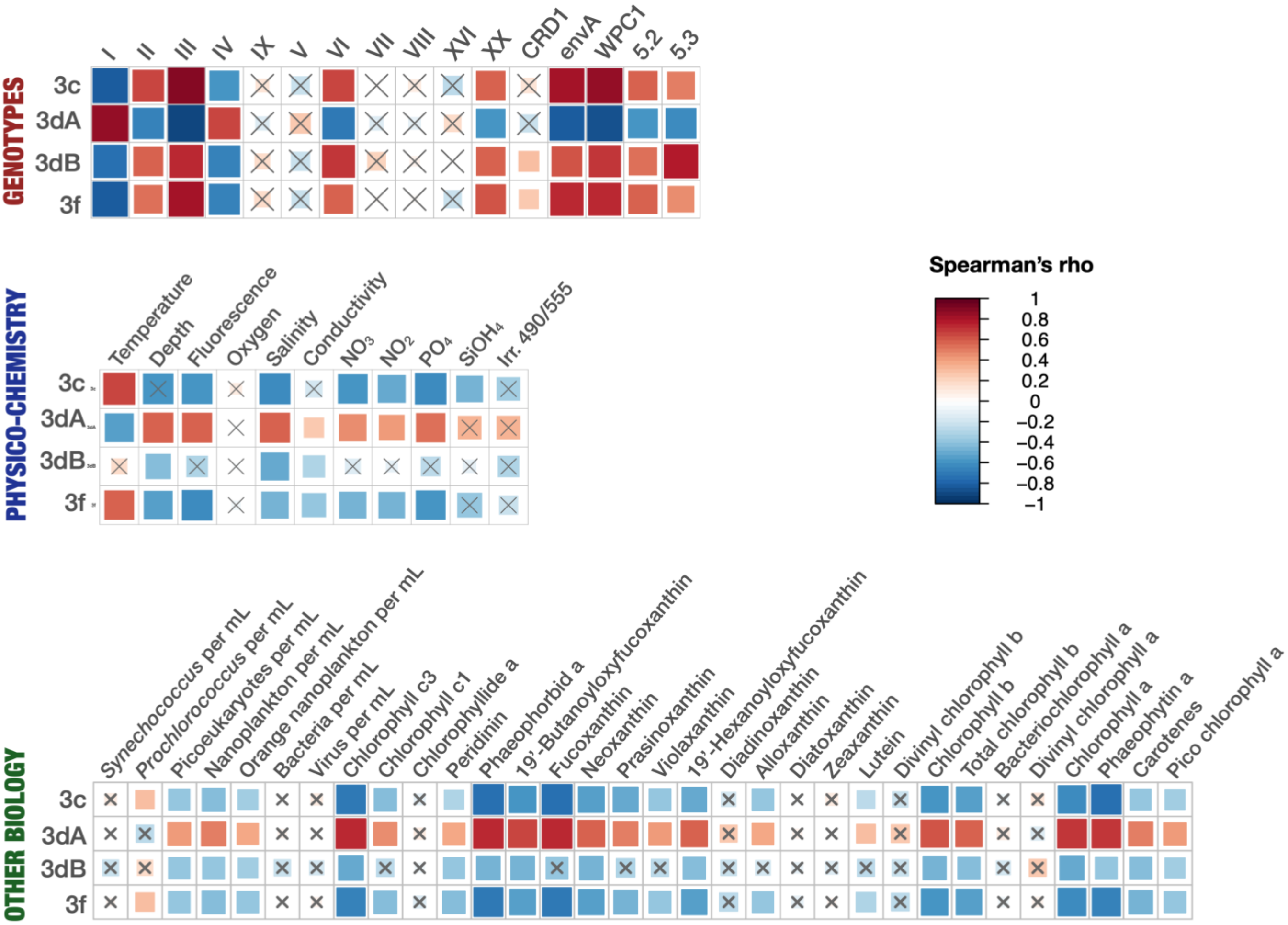


**Fig. S7: Correlation analysis between *Synechococcus* pigment types and all environmental and biological parameters measured at BOUSSOLE in the Mediterranean Sea.** Each row corresponds to a pigment type and each column to a parameter, separated in the three categories: *Synechococcus* genotypes, physico-chemical parameters and other biological parameters. The color scale indicates Spearman correlation coefficient between pairs of variables. Grey crosses indicate non-significant correlations (permutation-based test using 999 permutations, adjusted p-values > 0.05). Only pigment types and genotypes representing more than 5% of the *Synechococcus* reads in the whole dataset were represented. Abbreviations: Irr. 490/555, ratio of downward irradiance at 490 nm and 555 nm; Rrs. 490/555, ratio of remote sensing reflectance at 490 nm and 555 nm. See Dataset 1 for more details.

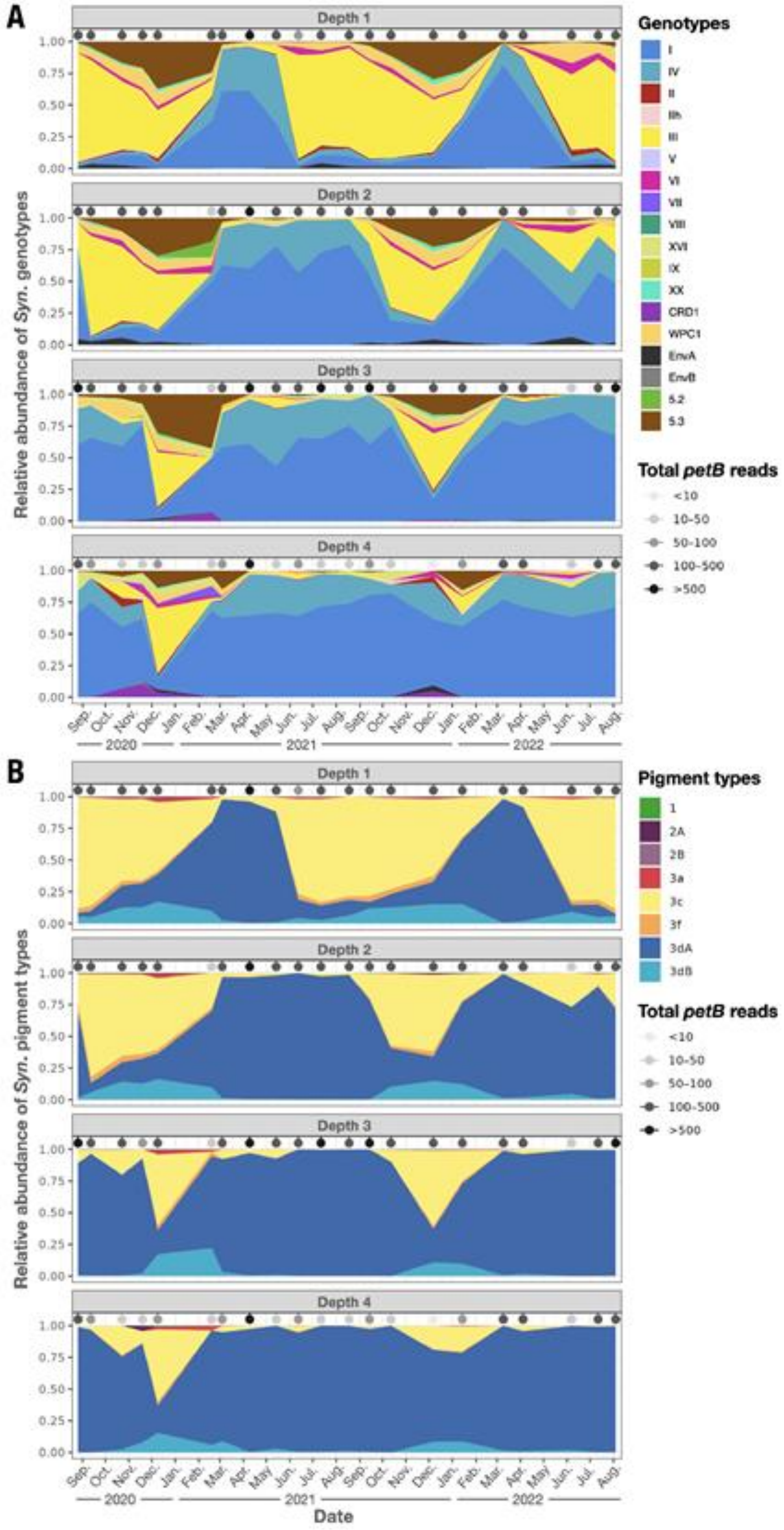


**Fig. S8: Seasonal variations in the abundances of *Synechococcus* genotypes (A) and pigment types (B) across depth at the Mediterranean station BOUSSOLE.** These abundances were computed from metagenomic reads recruitment on a set of reference gene markers (*petB* for genotypes; *cpcBA*, *mpeBA* and *mpeW* for pigment types). One area plot is plotted for each categorical depth, indicated in the grey bars. Depth 1 corresponds to the surface already shown in Fig. 1. Sampling points are indicated by black ticks and grey points on top of each area plot, with the intensity of grey reflecting the total amount of *petB* reads sequenced in each sample. Depth 1, surface; Depth 2, 20 m; Depth 3, Deep chlorophyll maximum; Depth 4, 60 m.

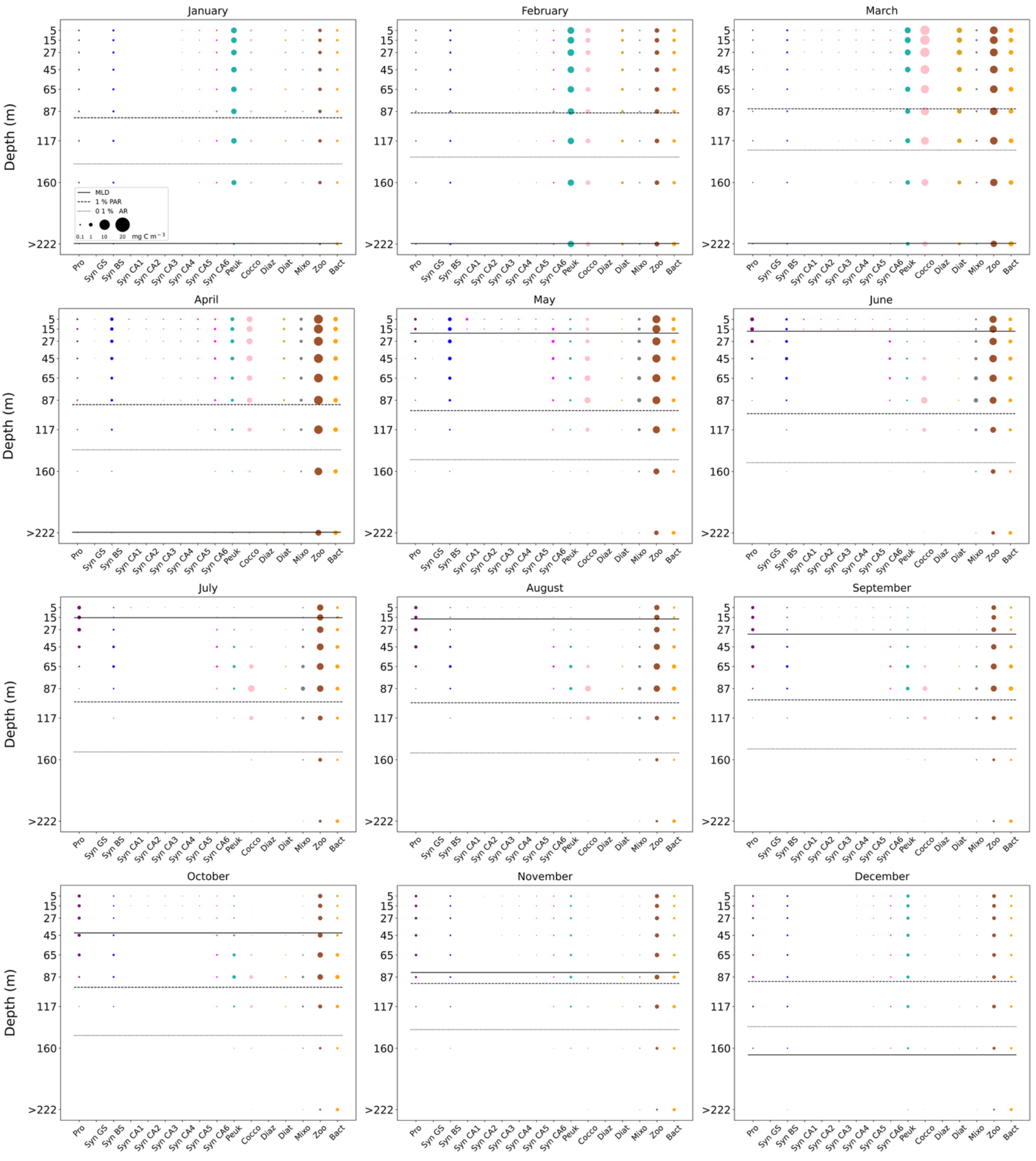


**Fig. S9 : Simulated biomass distribution of plankton functional groups (mg C.m$^{-3}$) at the closest proxy grid point (43.5°N, 7.5°E) to the BOUSSOLE station.** Bubble size is proportional to the biomass of the respective groups. The solid horizontal line represents the mixed layer depth, while the depths at which light intensity corresponds to 1% and 0.1% of surface PAR are indicated by the dashed and dotted lines, respectively.

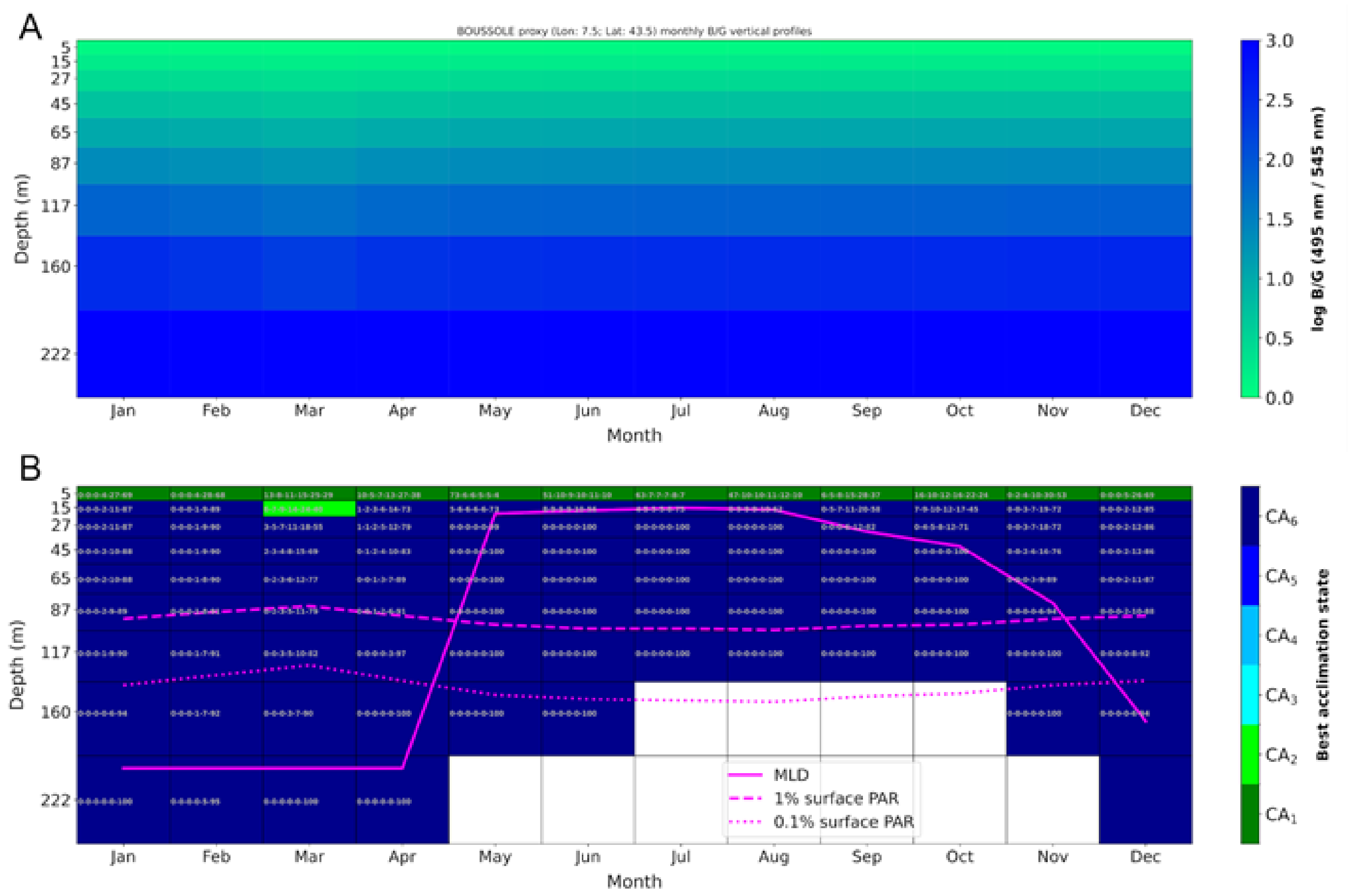


**Fig. S10 : Simulated vertical profiles of blue-to-green ratios for the closest grid point in the model domain, and monthly acclimation states of chromatic acclimaters**. **A** Monthly vertical B/G (495 nm/545 nm) profiles at the BOUSSOLE station (northwestern Mediterranean Sea). Columns represent monthly mean profiles (January–December), and rows correspond to discrete depth levels (5–222 m). Colors indicate the magnitude of the B/G ratio. **B** Monthly vertical profiles of the acclimation states for BOUSSOLE. The best acclimation state, indicated by the background color, was the state most efficient in harvesting the available light for a given portion of the water column and month of the year, representing the acclimation target for all CAs (direction of the acclimation process). The numbers within each depth box represent the relative contribution of each acclimation state to the total CA biomass, ordered from CA1 to CA6, with CA1 corresponding to the greenest acclimation state (i.e., lowest PUB:PEB) and CA6, the bluest acclimation state (i.e., highest PUB:PEB).
